# Specific gut microbiome members are associated with distinct immune markers in allogeneic hematopoietic stem cell transplantation

**DOI:** 10.1101/590893

**Authors:** Anna Cäcilia Masche, Katrine Kielsen, Malene Skovsted Cilieborg, Ole Lund, Susan Holmes, Frank M. Aarestrup, Klaus Gottlob Müller, Sünje Johanna Pamp

## Abstract

**Background:** Increasing evidence reveals the importance of the microbiome in health and disease and inseparable host-microbial dependencies. Host-microbe interactions are highly relevant in patients receiving allogeneic hematopoietic stem cell transplantation, (HSCT), i.e. a replacement of the cellular components of the patients’ immune system with that of a foreign donor. HSCT is employed as curative immunotherapy for a number of non-malignant and malignant hematologic conditions, including acute lymphoblastic leukemia. The procedure can be accompanied by severe side effects such as infections, acute graft-versus-host disease (aGvHD), and death. Here, we performed a longitudinal analysis of immunological markers, immune reconstitution and gut microbiota composition in relation to clinical outcomes in children undergoing HSCT. Such an analysis could reveal biomarkers, e.g. at the time point prior to HSCT, that in the future could be used to predict which patients are of high risk in relation to side effects and clinical outcomes and guide treatment strategies accordingly.

**Results:** In two multivariate analyses (sparse partial least squares regression and canonical correspondence analysis), we identified three consistent clusters: (1) High concentrations of the antimicrobial peptide human beta-defensin 2 (hBD2) prior to the transplantation in patients with high abundances of *Lactobacillaceae*, who later developed moderate or severe aGvHD and exhibited high mortality. (2) Rapid reconstitution of NK and B cells in patients with high abundances of obligate anaerobes such as *Ruminococcaceae*, who developed no or mild aGvHD and exhibited low mortality. (3) High inflammation, indicated by high levels of C-reactive protein, in patients with high abundances of facultative anaerobic bacteria such as *Enterobacteriaceae*. Furthermore, we observed that antibiotic treatment influenced the bacterial community state.

**Conclusions:** We identify multivariate associations between specific microbial taxa, host immune markers, immune cell reconstitution and clinical outcomes in relation to HSCT. Our findings encourage further investigations into establishing longitudinal surveillance of the intestinal microbiome and relevant immune markers, such as hBD2, in HSCT patients. Profiling of the microbiome may prove useful as a prognostic tool that could help identify patients at risk of poor immune reconstitution and adverse outcomes, such as aGvHD and death, upon HSCT, providing actionable information in guiding precision medicine.

## Background

The microbiome has gained increasing attention as a crucial contributor in the course of various diseases, and as target of treatment [1–3]. A number of complications in allogeneic hematopoietic stem cell transplantation (HSCT) have recently been associated with the gut microbiota [4, 5]. HSCT is a curative treatment for various hematologic diseases including malignancies, such as acute lymphoblastic leukemia (ALL), as well as non-malignant diseases, such as metabolic disorders and immune deficiency syndromes [6]. One goal of HSCT is achieving a beneficial graft-versus-leukemia (GvL) effect where donor-derived T lymphocytes and natural killer cells target leukemic cells in the recipient [7]. Prior to allogeneic HSCT, the patients undergo a preparative conditioning regimen involving combinations of chemotherapeutic agents and total body irradiation (TBI) [8] to eradicate leukemic cells and induce immunosuppression. Immunosuppressive treatment (both prior to and post HSCT) in the stem cell recipient prevents a graft-versus-host reaction caused by cytotoxic donor T lymphocytes that attack healthy cells in the recipient [8]. To limit infectious diseases due to immunosuppression, the patients are administered broad-spectrum antibacterial and antifungal compounds. Subsequently, the patients receive a stem cell graft originating from the bone marrow, peripheral blood or umbilical cord blood of a human leukocyte antigen (HLA)-matched sibling donor or an unrelated donor (i.e., allogeneic HSCT) [9, 10].

In patients undergoing HSCT, it has been previously observed that there are associations between the microbiome and clinical outcomes such as acute graft-versus-host disease (aGvHD) [5, 11, 12] and survival [4, 13]. GvHD after HSCT has been related to an expansion of the order *Lactobacillales*, especially *Enterococcus* spp. [11] and *Lactobacillus* spp. [14] and a loss of *Clostridiales* [14]. However, so far few studies have monitored the microbiome longitudinally [11, 14, 15]. This indicates the need for more detailed investigations that take temporal monitoring of both the host immune system and the microbiome into account.

Several markers of the host immune system, including inflammatory markers (such as C-reactive protein (CRP) and interleukin 6 (IL-6)) and markers of intestinal toxicity (such as plasma citrulline), have been studied in HSCT [16, 17]. Potential novel markers, such as antimicrobial peptides (AMPs), have also been proposed to be involved in outcomes after HSCT, for example in immunomodulation and regulation of microbial homeostasis [18]. AMPs, especially defensins such as human beta-defensin 2 and 3 (hBD2 and hBD3), have previously been found to play a role in some inflammatory diseases [19, 20]. However, to our knowledge, AMPs have not yet been employed as markers in the context of HSCT. An interesting research question remains: How are known and novel markers within the host immune system associated with each other or with changes in the microbiome? A better understanding about these associations would provide a more holistic insight into the underlying mechanisms affecting clinical outcomes after HSCT.

Another crucial factor impacting on complications and clinical outcomes after HSCT is immune reconstitution. Immune reconstitution involves the essential cellular components of the adaptive immune system, namely T and B cells, as well as key cellular components of the innate immune defense, namely natural killer (NK) cells, monocytes and neutrophils [21]. A microbial influence on immune cell differentiation has been observed previously, e.g., commensal *Clostridiales* were found to regulate Treg cell differentiation in the colon [22]. Therefore, an influence of the intestinal microbiome on immune reconstitution following HSCT is likely, but had not been investigated prior to this study.

Here, we monitor both host factors and the intestinal microbiome longitudinally. We included markers of inflammation (CRP and IL-6) and intestinal toxicity (plasma citrulline) as well as the antimicrobial peptides (hBD2 and 3), which in the following sections are collectively referred to as immune markers. To assess the prognostic potential of immune markers associated with gut microbial dynamics for immune reconstitution and clinical outcomes after HSCT (aGvHD and survival) in a holistic way, we implemented multivariate multi-table (also referred to as multi-way) approaches. This facilitated the integration of a variety of different factors that could influence the patients’ convalescence. We reveal distinct clusters of bacteria associated with sets of immune markers and clinical outcomes. Patients with rapid NK and B cell reconstitution that had no or mild aGvHD and low mortality exhibited high abundances of members belonging to the family of *Ruminocaccaceae*. In contrast, patients with moderate to severe aGvHD and high mortality showed high plasma concentrations of the antimicrobial peptide hBD2 already prior to HSCT. In these patients, we observed increased *Lactobacillaceae* abundances.

## Results

To assess associations between immune markers and gut microbial dynamics in the context of immune reconstitution and clinical outcomes after HSCT, we monitored 37 children over time undergoing allogeneic HSCT (Figure 1A, Additional file 1: Table S1). We gained insight into the patients’ immune reconstitution by determining T, B, NK, monocyte, and neutrophil cell counts in peripheral blood (Figure 1A, Additional file 1: Table S1). We measured C-reactive protein (CRP) and plasma interleukin 6 (IL-6) as markers of inflammation, and human beta-defensin 2 and 3 (hBD2 and 3) as markers for potential innate immune activation and systemic infection (Figure 1A, Additional file 1: Table S1). Plasma citrulline levels were measured as a marker of intestinal toxicity, being produced selectively by functioning enterocytes. To relate immune marker levels to members of the intestinal microbiota in patients undergoing HSCT, we characterized the longitudinal dynamics of the human intestinal microbiome in a subset of 30 patients by utilizing 16S rRNA gene profiling (Figure 1A).

**Figure 1.**
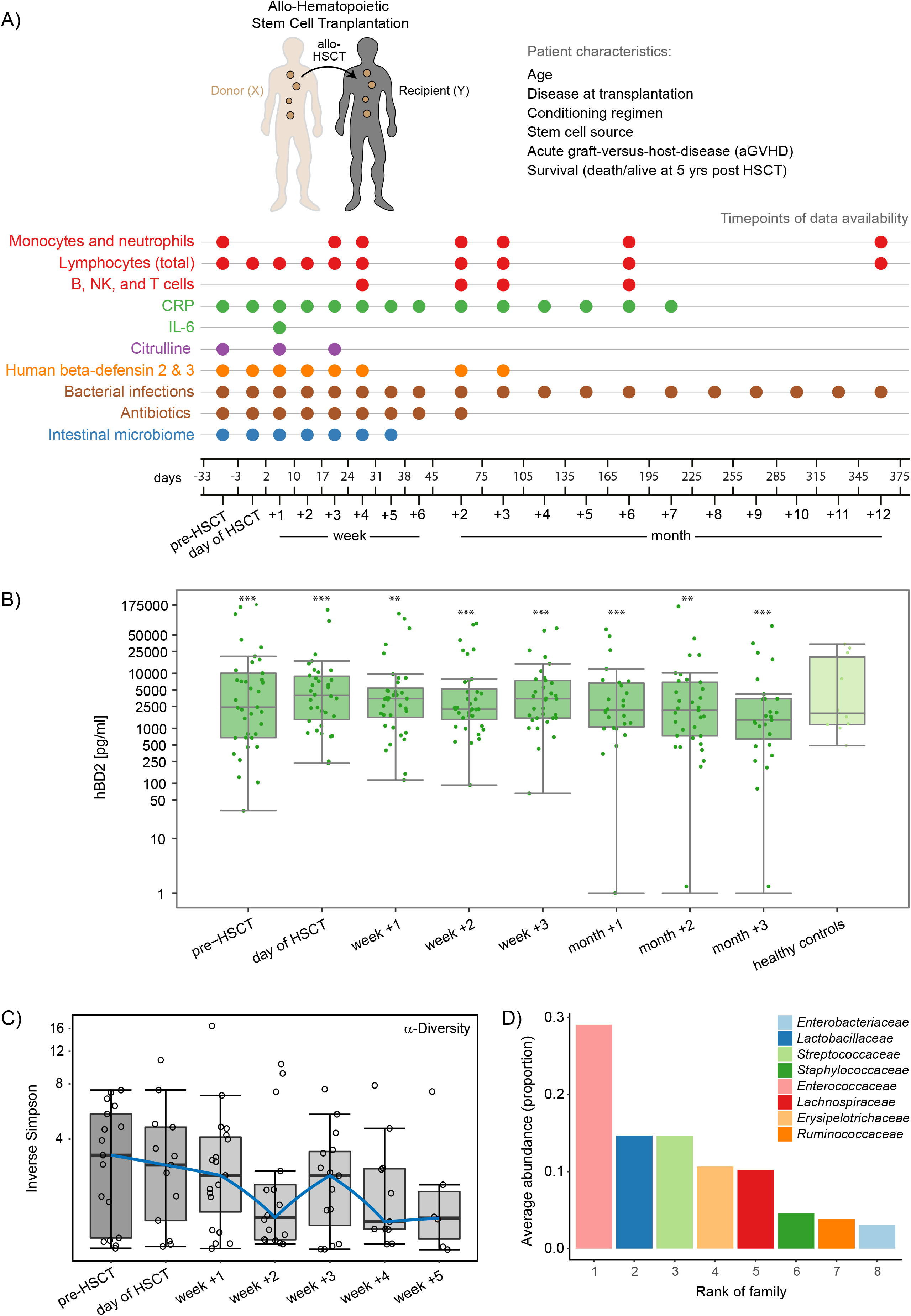
Monitoring of the host immune system and intestinal microbiome in hematopoietic stem cell transplantation (HSCT). (A) Study outline. A cohort of 37 pediatric recipients of allogeneic HSCT was monitored prior to HSCT, at the time point of HSCT, and post HSCT (median follow-up time 5.2 years). A range of patient characteristics as well as disease outcomes, immune markers, immune cell counts, and intestinal patterns of microbial community composition were recorded at the noted time points (up to 12 months post HSCT). See Table S1 for details regarding the patient characteristics. (B) Plasma hBD2 concentrations over time and in comparison to healthy controls. The Y-axis was log10-transformed for better visualization. Zeros were replaced with 1 to avoid undefined values on the log-transformed axis. Asterisks indicate whether the concentrations at each respective time point were significantly different from any of the other time points (showing the maximum significance level). (C) Bacterial alpha-diversity, measured by inverse Simpson index, of the intestinal microbiome shown with log10-transformed y-axis. (D) Rank abundance curve displaying the 8 most abundant taxonomic families in the dataset (comprising 98 fecal samples).

### Patient cohort and outcomes

At the time of HSCT, the 37 patients were on average 8.2 years old (age range 1.1 – 18.0 years). Twenty-five patients (68%) were diagnosed with at least one bacterial infection at median 75 days post HSCT (range: day −19 to +668). Twenty-six patients (70%) had no or mild aGvHD (grade 0-I) (Additional file 1: Table S1). Eleven patients (30%) developed moderate to severe aGvHD (grade II, III or IV) at median 18 days (range: day +9 to +45) after transplantation (Additional file 1: Table S1). Seven patients (19%) died during the follow-up period at median 266 days post HSCT (range: day +9 to +784) (5 relapse-related and 2 treatment-related deaths) (Additional file 1: Table S1). In total, six patients (16%) relapsed, four of which underwent a re-transplantation. All patients received antibiotics pre- and post-transplantation (Additional file 1: Table S1). Prophylactic trimethoprim-sulfamethoxazole was administered to all patients from day −7 until transplantation. During the period of neutropenia or latest from day −1, patients received prophylactic intravenous ceftazidime. In case of infections indicated by fever or microbial culture, ceftazidime was substituted by intravenous meropenem, vancomycin or other antibiotics, according to culture-based results.

### Temporal dynamics of immune markers and the intestinal microbiota in HSCT patients

Prior to assessing the interplay between clinical variables (i.e., immune markers, immune reconstitution, clinical outcomes) and the intestinal microbiota, we characterized these components separately. In order to provide an overview of changes in immune markers and immune cell counts after HSCT in our cohort (*n=37*), we assessed their temporal patterns (Figures 1B, 1C, and Additional file 2: Figure S1). Of note, these supplemental univariate analyses mainly serve the purpose of visually aiding our subsequent multivariate analyses approaches (Additional file 3: Figure S2). We characterized hBD2 for the first time in the context of HSCT by assessing plasma hBD2 concentrations over time from pre-HSCT to month +3 post HSCT in patients compared to healthy controls. The hBD2 concentration differed significantly between time points (*P* < 0.001, Kendall’s *W* = 0.6). It increased from pre-HSCT to the day of HSCT (*P* < 0.001), then slightly decreased in week +1 (*P* = 0.038) before increasing again in week +3 (*P* = 0.006). HBD2 decreased again in month +2 (*P* = 0.014) (Figure 1B). CRP levels differed significantly between time points (*P* < 0.001, Kendall’s *W* = 0.33). They were high pre-HSCT and until week +2, then decreasing significantly in week +3 (*P* < 0.001) with the lowest levels in weeks +4 to +6 (*P* < 0.001) (Additional file 2: Figure S1A). Median plasma citrulline levels were significantly different between time points (*P* < 0.001, Kendall’s *W* = 0.32). They decreased from pre-HSCT to week +1 (*P* < 0.001) and increased again in week +3 (*P* < 0.001) (Additional file 2: Figure S1A). B cell counts (Kendall’s *W* = 0.5) as well as CD4+ T cell counts (Kendall’s *W* = 0.46) increased steadily from month +1 to month +6 (*P* < 0.001) (Additional file 2: Figure S1B).

To gain insight into intestinal microbial dynamics before, at the time of, and after HSCT, we obtained a total of 97 fecal samples from a subcohort of 30 patients. Using 16S rRNA gene sequence analysis, we identified 239 operational taxonomic units (OTUs) (see Methods). Microbial alpha-diversity was lower at all time points post-HSCT compared to pre-HSCT (Figure 1B). The median inverse Simpson index decreased from 3.27 (range: 1.02 – 7.4) before HSCT to 2.89 (range: 1.04 – 10.77) on the day of transplantation and further to 2.03 (range: 1.0 – 16.51) post-HSCT (median of week +1 to +5). *Enterococcaceae* (*Firmicutes*) was the most abundant bacterial family observed, followed by other *Firmicutes*, such as three families from the class of *Bacilli* (*Lactobacillaceae*, *Streptococcaceae*, *Staphylococcaceae*), two families from the class of *Clostridia* (*Lachnospiraceae*, *Ruminococcaceae*), a family within the class of *Erysipelotrichia* (*Erysipelotrichaceae*) and a family within the class of gamma-*Proteobacteria* (*Enterobacteriaceae*) (Figure 1C).

### Associations between immune markers and immune cell reconstitution in HSCT patients

In order to identify patient baseline parameters and clinical outcomes (e.g. aGvHD, relapse, overall survival), as well as immune markers and immune cell types that might be important determinants in HSCT in relation to the microbiome, we performed variable assessment by permutational multivariate analysis of variance (adonis). The variables that were found to be significant (*P* ≤ 0.05), i.e., those that explained most variation in the microbial community distance matrix, were selected for subsequent analyses (Additional file 3: Figure S2, Additional file 4: Table S2). Of note, the occurrence of relapse as an indication of transplantation outcome was assessed but not found to be significant in adonis and was therefore not included in follow-up analyses. We then assessed associations of the selected immune markers and immune cells in the data set comprising 37 patients by determining Spearman’s rank correlations. The hBD2 concentrations pre-HSCT, on the day of HSCT and in weeks +1 and +2 post-HSCT were positively correlated with each other (ρ= 0.73 – 1, *P* < 0.001) (Figure 2A). NK cell counts in month +1 exhibited a positive correlation with total B cell counts (ρ= 0.64, *P* = 0.0046) and mature B cells counts (ρ= 0.62, *P* = 0.0114) in month +2. When we related immune cell reconstitution to outcomes, we observed significantly higher NK cell counts and total B cell counts in month +2 in patients with no or mild aGvHD (grade 0-I) compared with patients with moderate to severe aGvHD (grade II-IV) (Wilcoxon rank sum test, NK cells, *P* = 0.011; B cells, *P* < 0.001) (Figure 2B).

**Figure 2.**
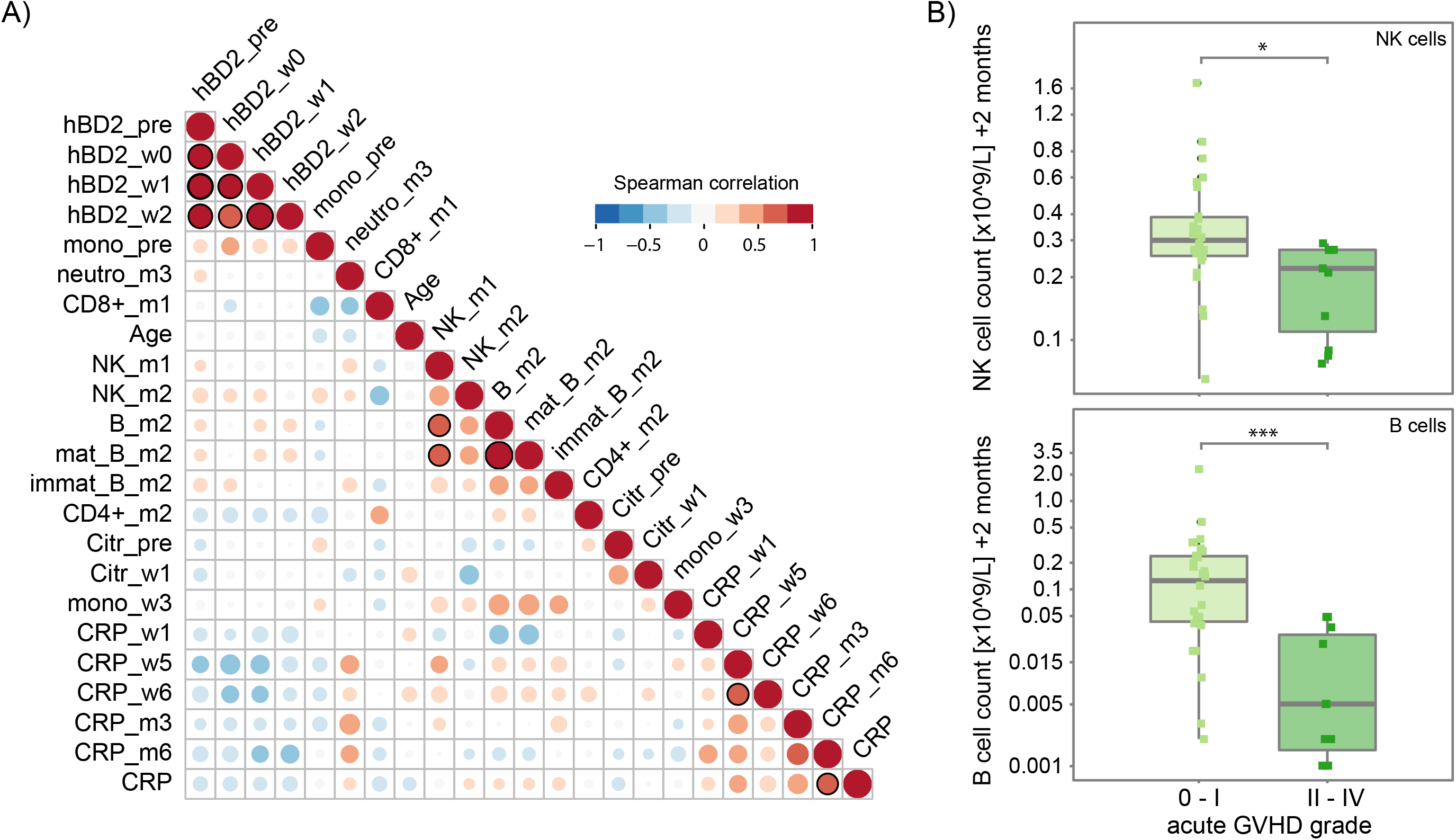
Correlations between immune markers, immune cell counts, and outcomes in patients undergoing HSCT. A) Pairwise Spearman’s correlation between immune markers and immune cell counts in HSCT patients (*n=37*) that were determined to be significant in a permutational multivariate analysis of variance using (microbial) distance matrices (adonis) (See Table S2). Positive and negative correlations are represented by red or blue circles, respectively, and the size of circles and intensity of color refer to the strength of the correlation. Correlations that are significant (*P* ≤ 0.05) are indicated by a black outline of the circle. (B) Natural killer (NK) and total B cell (mature and immature) reconstitution in month +2 with respect to the maximum acute GvHD (aGvHD) grade (0-I vs. II-IV) in HSCT patients (*n=37*). Abbreviations: hBD2_pre, hBD2_w0, hBD2_w1, hBD2_w2: plasma human beta-defensin 2 concentration pre-HSCT, on the day of HSCT, and in weeks +1 and +2, respectively; mono_pre, mono_w3: monocyte counts pre-HSCT and in week +3, respectively; neutro_m3: neutrophil count in month +3; CD8+_m1: CD8+ T cell counts in month +1; Age: Recipient age at time point of transplantation; NK_m1, NK_m2: Natural killer cell counts in months +1 and +2, respectively; B_m2, mat_B_m2, immat_B_m2: all, mature, and immature B cell counts in month +2; CD4+_m2: CD4+ T cell counts in month +2; Citr_pre, Citr_w1: plasma citrulline levels pre-HSCT and in week +1, respectively; CRP, CRP_w1, CRP_w5, CRP_w6, CRP_m3, CRP_m6: C-reactive protein levels at time points simultaneous to microbiome characterization, in weeks +1, +5, and +6, and in months +3 and +6, respectively. * *P* < 0.05, ** *P* < 0.01 and *** *P* < 0.001.

### High plasma hBD2 and monocytes prior to HSCT in patients with high *Lactobacillaceae*

To gain insight into how the selected immune markers and immune cell counts co-vary with gut microbial abundances in patients with distinct outcomes, we implemented two multivariate multi-table approaches for the subcohort (*n=30*), namely sparse partial least squares (sPLS) regression and canonical correspondence analysis (CCpnA). The sPLS regression models OTU abundances as predictors and clinical variables as response variables and explains the latter in an asymmetric (i.e. unidirectional) way. In contrast, the CCpnA assesses relationships between parameters of the immune system and microbiota bidirectionally. In the following paragraphs, the results of these two analyses are reported for one observed cluster at a time, respectively.

First, we performed sPLS regression to reveal multivariate correlation structures between immune markers, immune cell counts, and OTU abundances, modeling the latter as explanatory variables. The sPLS regression and subsequent hierarchical clustering suggested that the data separated into three clusters (Figure 3A). High monocyte counts and high plasma hBD2 concentration prior to HSCT, high patient age at the time of transplantation, and high abundances of *Lactobacillaceae* independent of time point contributed the most to the formation of cluster 1. Of note, monocyte counts and hBD2 were positively correlated with each other, in agreement with the correlation analysis above (Figure 2A). The *Lactobacillaceae* were represented by microaerophilic *Lactobacillus* sp. OTUs (e.g., AF413523.1, GU451064.1, KF029502.1) (Figure 3B, Additional file 5: Table S3). These OTUs exhibited high loading weights in sPLS dimension 2 (Figure 3C), indicating that they contributed strongly to the separation of clusters in dimension 2 (Figure 3A and 3B).

**Figure 3.**
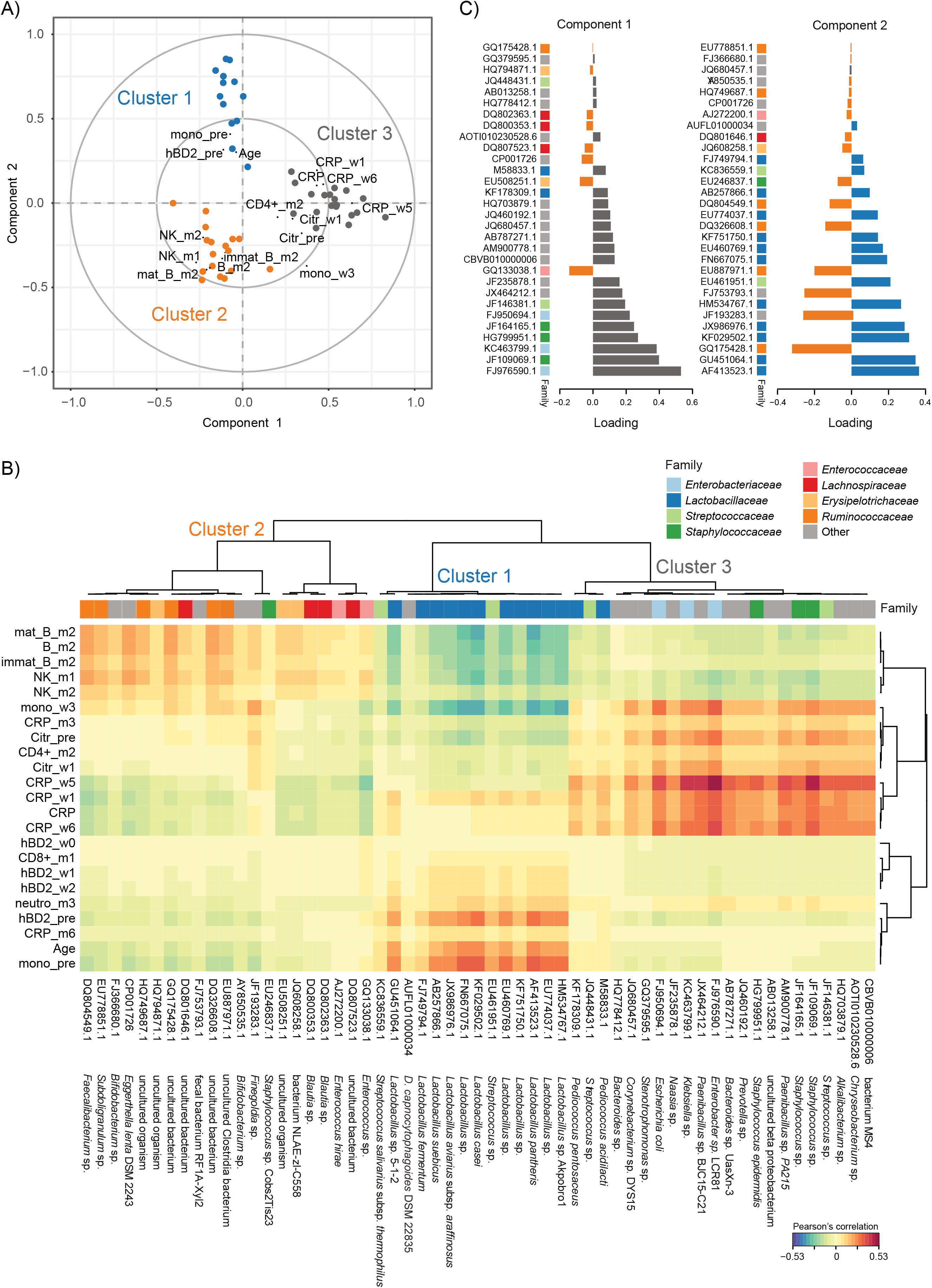
Sparse partial least squares (sPLS) regression of immune parameters and intestinal bacterial taxa during HSCT. (A) Correlation circle plot for the first two sPLS dimensions with correlations displayed for >0.2/<−0.2. The two grey circles indicate correlation coefficient radii at 0.5 and 1.0. Bacterial operational taxonomic units (OTUs) are displayed as circles, and are colored according to the cluster they are affiliated with (Cluster 1: blue; Cluster 2: orange; Cluster 3: grey). Variables projected closely to each other are positively correlated. Variables projected diametrically opposite from each other are negatively correlated. Variables situated perpendicularly to each other are not correlated. (B) Clustered image map (CIM) of the first two sPLS dimensions, displaying pairwise correlations between OTUs (bottom) and clinical variables (left). Red and blue indicate positive and negative correlations, respectively. Hierarchical clustering (clustering method: complete linkage, distance method: Pearson’s correlation) was performed within the mixOmics *cim()* function based on the sPLS regression model. An overview of the OTU abundances in the individual samples is provided in Figure S3, and a list of the individual OTUs and their cluster-affiliation is provided in Table S3. (C) Loading plots of OTUs with maximum contributions on the first (left) and second (right) component, respectively. The bars are coloured according to the cluster they are affiliated with. The family-affiliation for each respective OTU is indicated by color (for legend see B). Abbreviations of variables are the same as in Figure 2.

Second, we applied CCpnA to model the canonical relationships between OTU abundances and clinical variables through the construction of common “latent” variables. The CCpnA confirmed the separation of the data into three clusters as observed in the sPLS regression (Figures 3A, 4, and Additional file 6: Figure S3A), including the clustering of OTUs. In addition, the CCpnA facilitated the inclusion of categorical variables, such as the patients’ baseline parameters (e.g., recipient sex, donor type) and clinical endpoints (aGvHD grade, overall survival). Because CCpnA is an unsupervised method and upheld the results of the sPLS regression, it provides confidence in the cluster findings.

Cluster 1 in the CCpnA seemed to include patients who developed moderate to severe aGvHD (grade II - IV) and who died. As suggested by both, sPLS and CCpnA, these patients exhibited high levels of plasma hBD2 before HSCT (and in week +1 and +2) and high monocyte counts before HSCT. OTUs within this CCpnA cluster predominantly included members of the family *Lactobacillaceae* and were most abundant in fecal samples of these patients (Figure 4 and Additional file 7: Figure S4). OTUs that were assigned to cluster 1 by the sPLS-based hierarchical clustering were congruently associated with the same clinical variables in the CCpnA (Figures 3B and 4).

**Figure 4.**
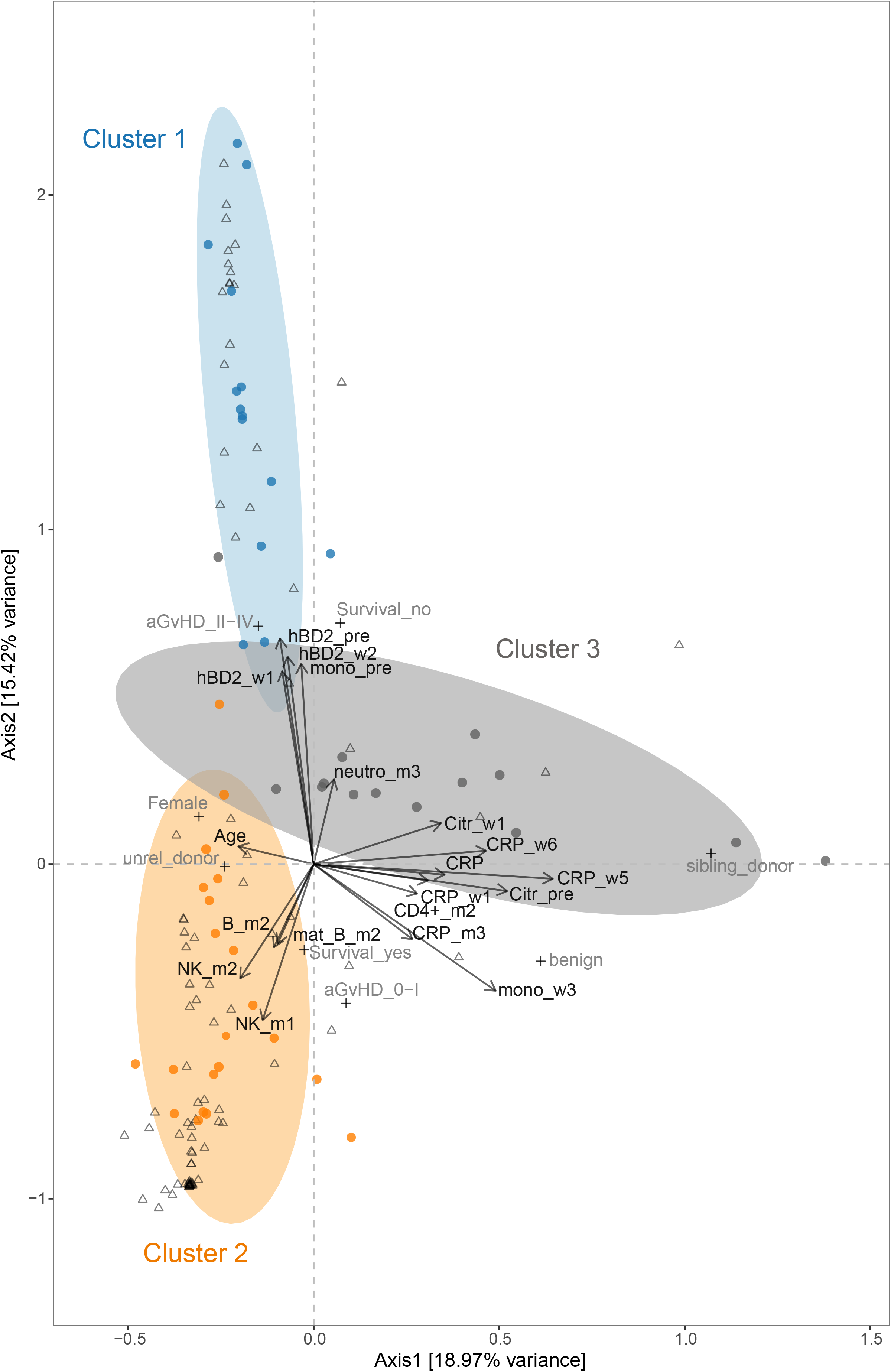
Canonical correspondence analysis (CCpnA) of immune parameters and intestinal bacterial taxa in patients undergoing HSCT. Triplot showing dimension 1 and 2 of the CCpnA that included continuous clinical variables (arrows), categorical variables (+), and OTUs (circles). Samples are depicted as triangles. OTUs with a correlation of >0.2/<−0.2 in the sPLS analysis were included in the CCpnA model. Only the variables and OTUs with a score >0.2/<−0.2 in at least one CCpnA dimension are shown. The OTUs in the CCpnA plot are colored according to the cluster they were affiliated with in the sPLS-based hierarchical clustering analysis, and the ellipses present an 80% confidence interval, assuming normal distribution. For visualization purposes, this plot is a section focussing on the categorical and continuous variables contributing to the ordination. The full size version of the CCpnA triplot, including all samples and OTUs, is presented in Figure S2A. Abbreviations of variables are the same as in Figure 2.

### Temporal patterns of the *Lactobacillaceae*-dominated community state type

Upon revealing an association between high plasma hBD2 concentrations and monocyte counts prior to HSCT, moderate to severe aGvHD, and high mortality with high abundances of *Lactobacillaceae* in multivariate analyses, we assessed in more detail the importance of longitudinal changes in these components. We implemented an additional approach to identify distinct bacterial community patterns by employing partitioning around medoid (PAM) clustering (see Methods). In this analysis, we detected four community state types (CSTs). Dominating taxa, similar to those identified by the sPLS-based hierarchical clustering, were revealed in the CSTs, e.g., *Lactobacillaceae* members dominated CST 1 (Additional file 5: Table S3 and Additional file 8: Figure S5). We then used this information to examine temporal community state changes in individual patients, i.e., their transitions between CSTs over the course of time (Figure 5). Based on clinical outcomes, the patients can be divided into four groups: 1. Patients who developed no or mild aGvHD (aGvHD grade 0-I) and survived compared to 2. Patients who died, and 3. Patients who developed moderate to severe aGvHD (grade II-IV) and survived compared to 4. Patients who died (Figure 5). We observed that 6 out of 8 patients (75%) with moderate to severe aGvHD (groups 3 and 4) harboured the *Lactobacillaceae*–dominated CST 1 at least once during the monitored period (1x: *n=2*, 2x: *n=2*, 3x: *n=2*) (Figure 5). In comparison, only 5 out of 22 patients (23%) with aGvHD grade 0-I (groups 1 and 2) carried CST 1 one or two times (Figure 5). Interestingly, high abundances of *Lactobacillaceae* in patients with aGvHD grade II-IV (groups 3 and 4) occurred predominantly at late time points (week +1 and later) (Figure 5 and Additional file 7: Figure S4).

**Figure 5.**
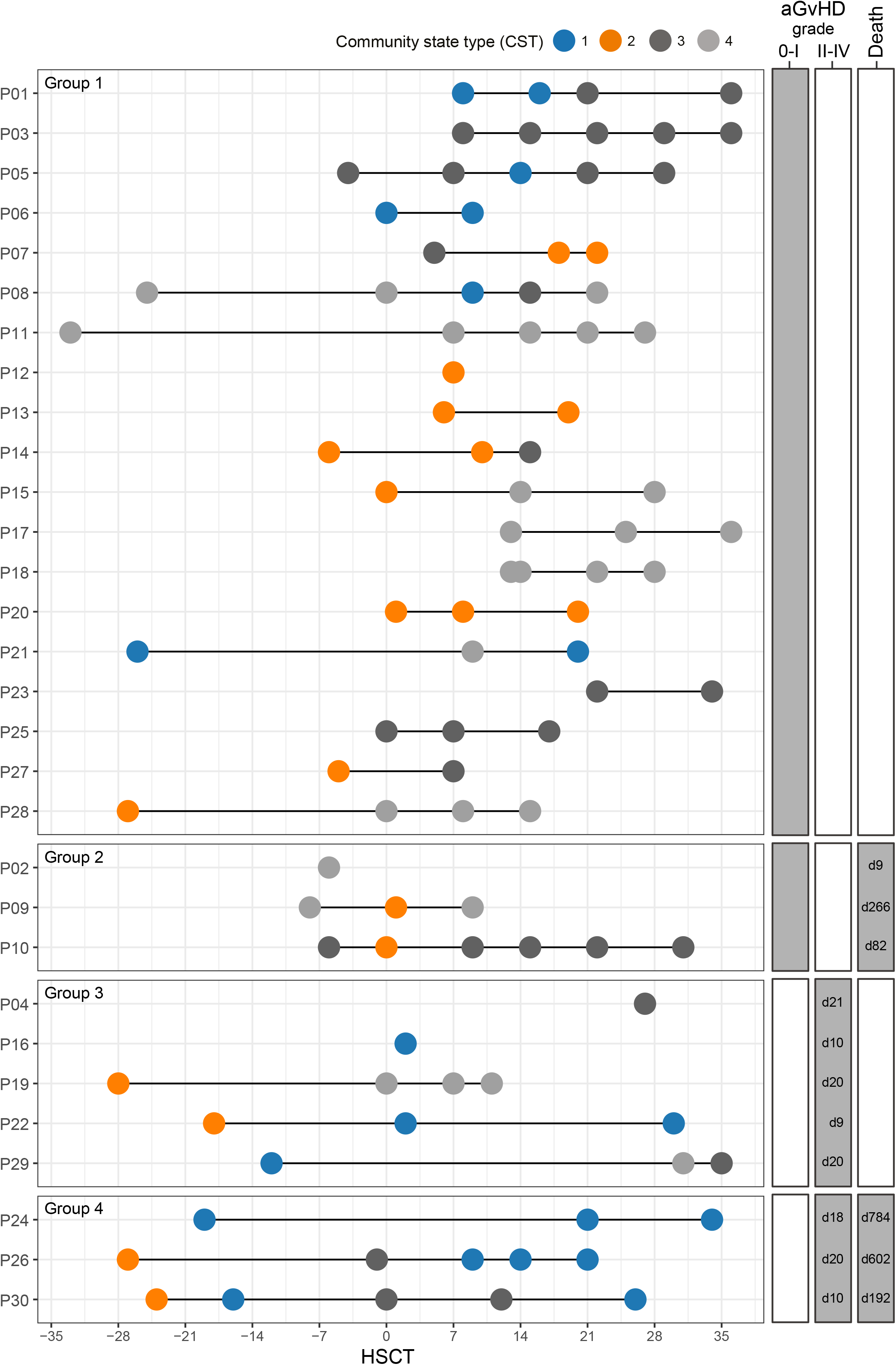
Bacterial community state types over time in patients undergoing allogeneic HSCT. Patients are grouped into four outcome groups: 1. Patients who developed no or mild aGvHD (grade 0-I) and survived vs. 2. Patients who did not survive; 3. Patients who developed moderate to severe aGvHD (grade II-IV) and survived vs. 4. Patients who did not survive. The day of commencement of aGVHD grade II-IV and the day of death post HSCT are displayed to the right. Patients with moderate to severe aGvHD (groups 3 and 4) most often harbored the *Lactobacillaceae-*dominated community state type 1 (CST 1), especially at late time points. CST2, dominated by *Ruminococcaceae* and *Lachnospiraceae*, did not persist after HSCT in any of the patients in groups 3 and 4. A detailed overview of the CSTs is provided in Figure S4, and information about individual OTUs and their cluster-affiliation is provided in Table S3.

### Temporal association of *Lactobacillaceae* with aGvHD and immune markers

To relate temporal changes in immune markers to those in the bacterial community composition in patients with aGvHD grade II-IV who died (group 4), we assessed their individual longitudinal profiles (Figure 6). In all three patients (P24, P26, P30), a large expansion of *Lactobacillaceae* abundances after the onset of aGvHD was observed. The average relative abundance of *Lactobacillaceae* after aGvHD onset was 72.92% (range 0.22 – 97.04%), as compared to before aGvHD (average 9.88%, range 0.25 – 37.83%) (Figure 6). Furthermore, bacterial alpha diversity was lower at the time point after aGvHD onset compared to the time point before (Figure 6). All three patients were treated with antibiotics for different durations between these two time points and prior to the time point before aGvHD onset.

**Figure 6.**
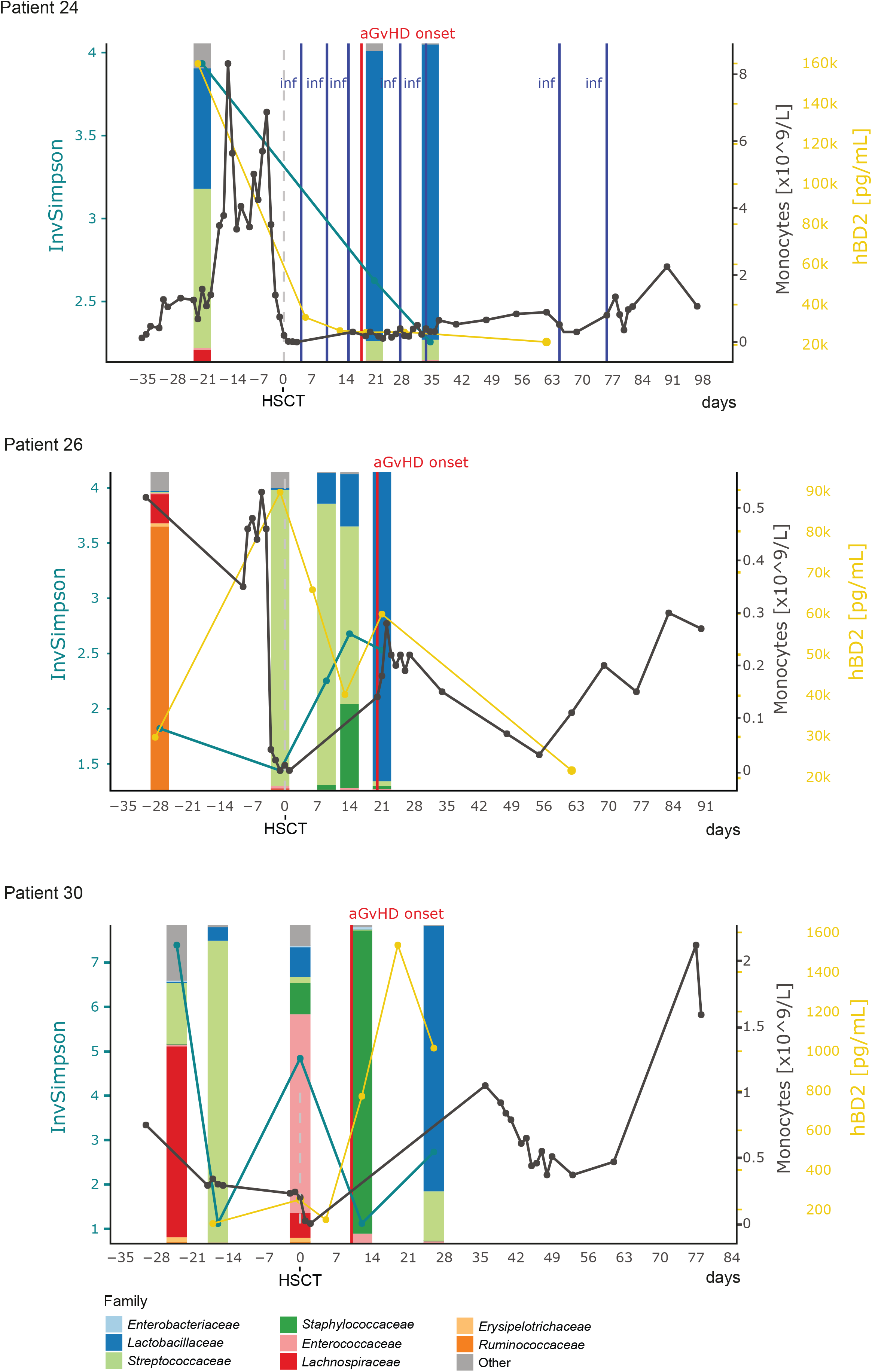
Longitudinal profiles of microbial community composition and immune markers in non-survivors with aGvHD. Abundances of *Lactobacillaceae* increased predominantly after aGvHD onset in patients who died during the follow-up period. Patient P24 developed chronic GvHD on day +187, relapsed on day +548 and died on day +784 due to graft rejection after re-transplantation. Patient P26 died on day +602 after a relapse on day +442 followed by re-transplantation on day +518. Patient P30 died on day +192 after relapse on day +77. Patient P26 and P30 had no reported bacterial infections during the depicted period. Abbreviations: InvSimpson, inverse Simpson diversity index; hBD2, human beta-defensin 2; aGvHD, acute graft-versus-host-disease; inf, bacterial infection.

In agreement with the results of the multivariate analyses, two of the patients (P24 and P26) exhibited between 1.95 and 14.56 times higher plasma hBD2 concentrations for all measured time points compared with the average of the whole data set (average plasma hBD2 concentration: 10,983.9 pg/ml, range: 0-177,400.28 pg/ml). The plasma hBD2 concentration was the highest at the time of HSCT in patient P26 and before HSCT in patient P24 (Figure 6). Of note, patient P24 received corticosteroid-based GvHD prophylaxis prior to HSCT and both patients received anthymocyte globulin as part of their conditioning regimen. These two patients had underlying malignant diseases and their pre-HSCT CRP levels, providing insight into underlying inflammation, were 1.6 times lower and 1.29 times higher compared with the average of the whole data set on that time point (average plasma CRP concentration before HSCT: 19.21 mg/L:, range: 1 – 60.36 mg/L), respectively. Monocyte counts before HSCT, i.e., recipient-derived cells, were 6.25 times higher in patient P24 compared with the average of the whole data set (average monocyte count before HSCT: 0.45 × 10^9^/L, range: 0.08 – 2.83 × 10^9^/L) and close to average in patient P26. In both patients, monocyte counts were higher before (i.e., recipient-derived cells) than after HSCT (i.e., donor-derived cells) (Figure 6).

In contrast to the *Lactobacillaceae* expansion after aGvHD onset in group 4, we observed high *Lactobacillaceae* abundances already before aGvHD onset in three survivors (P16, P22 and P29) who developed aGvHD grade II-IV (group 3) (Additional file 9: Figure S6). The average relative abundance of *Lactobacillaceae* before aGvHD onset was 53.18% (range 0.35 – 98.86%), as compared to after aGvHD (average 18.25%, range 0 – 54.78%). However, these observations were limited by a small number of patients per outcome group (groups 3 and 4) (Figure 5) and therefore cannot serve to provide statistical evidence.

### High NK and B cells and no or mild aGvHD in patients with high obligate anaerobes such as *Ruminococcaceae*

The association between NK and total B cell counts after HSCT observed in the Spearman’s correlation analysis (Figure 2) was supported by the multivariate analyses (Figures 3 and 4). In the sPLS regression, cluster 2 included high NK cell counts in months +1 and +2, as well as high immature, mature, and total B cell counts in month +2 (Figure 3A). Furthermore, NK and B cell counts were positively correlated with OTUs found in cluster 2 (Figure 3A and B), in particular with members of the family *Ruminococcacea* (Order: *Clostridiales*) (Figure 3B, Additional file 5: Table S3). Strong positive correlations between mature and total B cell counts in month +2 and NK cell counts in month +1 were observed, for example, with *Faecalibacterium sp*. (DQ804549.1) (Figures 3B and 3C, Additional file 5: Table S3). Additionally, we observed positive associations of these immune cells with *Lachnospiraceae* (Order: *Clostridiales*), among those two *Blautia* spp. (DQ800353.1 and DQ802363.1) (Figures 3B and 3C, Additional file 5: Table S3).

In support of the sPLS regression, the CCpnA also revealed high NK and B cell counts in cluster 2 in months +1 and +2. Based on the CCpnA, we found that NK and B cell reconstitution was associated with certain disease outcomes. For example, patients who had no or mild aGvHD and survived were predominantly represented in cluster 2. Cluster 2 included OTUs mainly belonging to *Ruminococcaceae* and *Lachnospiraceae*, exhibiting their highest abundances in patient samples associated with this cluster. OTUs with the highest scores in dimension 2 predominantly belonged to the family of *Ruminococcacea* and matched the sPLS-based hierarchical cluster assignment (Figure 4).

As all patients received antibiotic treatment prior to and post transplantation, we examined the potential effect of antibiotics on the intestinal bacterial community composition. A trending influence of the vancomycin treatment was revealed by adonis analysis (*P* = 0.055). Using a CCpnA model that also included information about antibiotic treatments, we found that patients exhibited higher abundances of *Ruminococcaceae* and *Lachnospiraceae* and lower abundances of *Enterobacteriaceae* at time points without the vancomycin treatment compared to with vancomycin (Additional file 6: Figure S3B). The same pattern was observed for treatment with ciprofloxacin (Additional file 6: Figure S3B).

Community state typing also revealed a state type dominated by *Ruminococcaceae* and *Lachnospiraceae* family members (CST 2) (Additional file 8: Figure S5). A total of 10 out of 22 patients (45%) with no or mild aGvHD were assigned to CST 2 at least at one time point (1x: *n=6*, 2x: *n=3*, 3x: *n=1*) (Figure 5). Half of the patients (4 out of 8) with moderate or severe aGvHD were also assigned to CST 2 once, but only at the time point before transplantation (Figure 5). That is, a *Ruminococcaceae-* and *Lachnospiraceae-*dominated community only persisted in patients with no or mild aGvHD.

### Persistence of *Enterococcaceae*-dominated community state type

A subcluster of cluster 2 comprised two facultative anaerobe *Enterococcus* spp. (GQ1330038.1 and AJ272200.1), exhibiting positive correlations with high NK and B cell counts (Figure 3B) and contributing to the separation of the clusters (Figure 3C). *Enterococcaceae* was the most abundant family in the overall study population (Figure 1D), and community state typing revealed a state type predominantly characterized by *Enterococcaceae* (CST 4, Additional file 8: Figure S5). A total of 9 out of 22 patients (41%) with no or mild aGvHD were assigned to CST 4 at least at one time point (1x: *n=2*, 2x: *n=2*, ≥3x: *n=5*) and this *Enterococcaceae*-dominated CST often persisted in the patient over time (Figure 5). A quarter of the patients (2 out of 8, both survivors) with moderate or severe aGvHD were assigned to CST 4 at least at one time point on the day of or post-HSCT (Figure 5).

### High inflammation in patients with high facultative anaerobic bacteria

In multivariate analyses, OTUs belonging to facultative anaerobic *Enterobacteriaceae* and *Staphylococcaceae* were characteristic for sPLS cluster 3 (Figures 3A and 3B, Additional file 5: Table S3). Cluster 3 was further comprised of high plasma citrulline concentrations pre-HSCT (before conditioning) and week +1, high monocyte counts in week +3, high CD4+ T cell counts in month +2, high CRP levels, particularly in weeks +1, +5, and +6 (Figures 3A and 3B). The projection of the variables in the sPLS suggested a weak positive correlation between these clinical variables in dimension 1 (Figure 3A). However, a weak negative association between CRP levels and monocytes (week +3) as well as citrulline and CD4+ T cell counts is indicated in dimension 2 (Figure 3A), which is in agreement with some results of the Spearman’s rank correlation tests (Figure 2).

The clinical variables in cluster 3, in particular high CRP levels post HSCT, exhibited positive correlations with the OTUs predominantly affiliated with *Proteobacteria* (e.g., *Enterobacteriaceae*), *Bacteroidetes*, and *Staphylococcus* spp. (Figure 3B, Additional file 5: Table S3). The strongest positive correlations occurred with several facultative anaerobe bacteria, e.g., *Enterobacter* sp. LCR81 (FJ976590.1), *Escherichia coli* (FJ950694.1) and *Staphylococcus* sp. (JF109069.1). The CCpnA supported observations from the sPLS regression regarding cluster 3, but indicated that they were only represented by a few patient samples (Figure 4 and Additional file 6: Figure S3A). These samples were characterized by high CRP levels, especially in week +1, +5, and +6. High CRP levels at these time points were not associated with aGvHD grade, i.e. they were not higher in patients with either aGvHD grade 0-I, or grade II-IV.OTUs of cluster 3, i.e., members of *Enterobacteriaceae and Staphylococcaceae*, exhibited their highest abundances in samples of this cluster (Figure 4 and Additional file 6: Figure S3A). Of note, patients represented in cluster 3 were younger compared to those in the other two clusters (Figure 4).

## Discussion

The human immune system and host-associated microorganisms are closely interlinked and play central roles in health and disease. The underlying components and mechanisms facilitating interactions between the immune system and microorganisms are however not completely understood. Understanding these associations is particularly relevant in patients that receive components of a “foreign” immune system, such as in patients undergoing allogeneic HSCT. Because both the patients’ immune system and microbiome appears severely affected, they potentially jointly impact on clinical outcomes. Here, we perform an integrated analysis of immune markers, immune reconstitution data, clinical outcomes, and microbiota, and we provide evidence for the association between specific microbial taxa, host immune markers, immune cells and clinical outcomes.

We observed that a predominance of *Clostridiales*, represented by *Ruminococcaceae* and *Lachnospiraceae*, in the intestine did not persist after transplantation in patients who either developed aGvHD grade II-IV, died, or both. *Clostridiales* are common colonizers of the healthy distal gut [23], and their loss is associated with microbial community disruption, reduced diversity, and GvHD [14]. A low diversity before conditioning and at the time of engraftment is associated with increased mortality [4, 5], in line with our findings.

In addition, in our multivariate analyses we observed high plasma hBD2 levels before HSCT and in weeks +1 and +2 post HSCT in patients who developed aGvHD grade II-IV, died, or both. The reason for the high hBD2 levels before HSCT in patients with increased mortality and GvHD is unknown at present. One could speculate that it relates to a higher burden of inflammation before and during the first two weeks post-transplantation. This was also reflected in higher counts of (recipient-derived) monocytes pre-HSCT that, by secreting hBD2, might contribute to an inflammatory reaction and thereby a potentially higher risk of aGvHD after HSCT. High hBD2 concentrations in weeks +1 and +2 could be an indicator for an innate immune reaction involving donor-derived cells, for example, against opportunistic pathogens that may have translocated to the bloodstream [24]. This might have been promoted by the decrease in *Ruminococcaceae* and *Lachnospiraceae* abundances in these patients after HSCT, indicating a microbial community disruption. Plasma hBD2 concentrations were characterized for the first time here in HSCT patients, and our findings emphasize the importance for further investigations to exploit the potential of hBD2 as a novel candidate marker for outcomes after HSCT. Importantly, our data suggested that differences in hBD2 secretion levels between patients were highly dependent on both, the microbial community composition and the time point relative to HSCT. We suggest that further investigations take into account variations of microbial community patterns and temporal changes when assessing hBD2 in the HSCT context. Moreover, these findings may be refined with a sampling time point homogeneity that is higher than what we provide in this study.

An interesting observation was that high pre-HSCT hBD2 levels and monocyte counts were associated with high abundance of *Lactobacillaceae* independent of time point. Probiotic *Lactobacillus* spp. have previously been shown to enhance hBD2-secretion in immune cells, thereby contributing to the innate immune defense [25, 26]. This mechanism could play a protective role during blood stream infections by opportunistic pathogens in HSCT patients. The increase of *Lactobacillaceae* abundances, particularly after the onset of aGvHD in patients who died, may be explained by a previously proposed compensatory mechanism to reduce aGvHD severity after onset in mice and humans [14]. For instance, high abundances of *Lactobacillaceae* could indicate homeostasis in the gut microbiome of children and thereby prevent inflammation caused by opportunistic pathogen expansion [27, 28]. Reduced intestinal inflammation might then benefit the outcome of aGvHD. This might however also lead to a less effective graft-versus-leukemia (GvL) effect as suggested by the finding that all patients, for which we observed moderate to severe aGvHD and subsequent increased *Lactobacillaceae* post HSCT, overcame aGvHD, but died following a relapse. Only a few patients in our study represented this combination of outcomes (moderate to severe aGvHD with subsequent increase of *Lactobacillaceae* and death), therefore no significant conclusion can be drawn at this point and further studies are needed to address these observations in more detail. We provide a discussion on survival following high *Lactobacillaceae* abundances prior to the onset of aGvHD in Additional file 10: Supplementary discussion.

In contrast to patients with moderate to severe aGvHD, we observed a higher overall survival in patients with no or mild aGvHD in multivariate analyses. The latter patients also had increased numbers of NK and B cells. In agreement with our study, previous studies have shown slower NK cell reconstitution after HSCT in patients with moderate to severe aGvHD compared to those with no or mild aGvHD [29]. Moreover, our results are in agreement with the previously described association of low NK cell numbers after HSCT and reduced overall survival [30], in line with NK cells’ crucial role in the GvL effect [7]. A poor recovery of B cells has been found to pose an increased risk of late infections [31], which might be an explanation for the association of high B cell counts and lower survival. The lower B cell numbers we observed in patients with more severe aGvHD could partially be a consequence of aGvHD or the treatment with corticosteroids, which is known to reduce the number of B cell precursors [21, 29]. However, based on our findings, the intestinal microbiota could also play a contributing role, because faster NK and B cell reconstitution as well as lower aGvHD severity, and higher overall survival were associated with high abundances of obligate anaerobes belonging to *Ruminococcaceae* and *Lachnospiraceae*. Indeed, decreased abundances of these bacterial families have previously been associated with an overall microbial disruption and decreased diversity [5, 15, 32]. In line with our observations, reduced microbial diversity was shown to contribute to lower survival [4] and a reduction of *Clostridiales* was observed in patients presenting aGvHD [33]. Furthermore, in melanoma patients, a low diversity and decreased *Ruminococcaceae* abundance were associated with a poor response to immunotherapy [34]. A potential explanation for this association could be that members of the order *Clostridiales* can downregulate inflammation and might thereby prevent aGvHD. Anti-inflammatory components produced by *Clostridiales* include, for example, urinary 3-indoxyl sulfate (3-IS) [35] and butyrate [36]. Acute GvHD might also reinforce microbial community disruption, as it is known to be accompanied by a reduction of Paneth cell numbers in the intestine. Paneth cells are secretors of α-defensins, important modulators of gut microbial homeostasis [37].

In contrast to the patient group with high abundances of obligate anaerobic bacteria (e.g., *Ruminococcaceae* and *Lachnospiraceae*), patients with high abundances of facultative anaerobic bacteria (e.g., *Enterobacteriaceae*, *Staphylococcus* spp., and *Streptococcus* spp.), showed slow NK and B cell reconstitution. An increase in facultative anaerobic bacteria has previously been observed in the gut microbiome of HSCT recipients before the start of conditioning compared with donors [5]. Here, we additionally observed high levels of C-reactive protein (CRP), indicating high inflammation in these patients. A possible explanation for this association might involve the shift to microbial growth conditions that favor facultative anaerobic bacteria during intestinal inflammation, such as an increased availability of oxygen caused by inflammatory products [38]. Our multivariate analyses indicated that the patients in which we observed these associations were younger compared to the rest of the cohort. Therefore, one could speculate that an immature intestinal microbiome might exhibit a higher susceptibility to opportunistic growth of facultative anaerobic bacteria. Further investigations will have to elucidate this relation and its potential consequences for adjustments of monitoring and treatment by age.

We provide a discussion on our findings regarding associations of adverse outcomes with *Enterococcus* compared with previous studies in Additional file 10: Supplementary discussion.

Antimicrobial treatment of patients can significantly affect the gut microbiota, especially in children [39]. Early use of antibiotics in general has been found to reduce *Clostridiales* in the intestine of HSCT patients [40]. Furthermore, a link of high total amounts of antibiotic in GvHD development has been observed in pediatric stem cell recipients [41]. However, to our knowledge, the effects of specific antibiotics on the intestinal microbiome especially in pediatric HSCT patients have not yet been elucidated in detail. Here, we identified a number of specific antibiotics, including vancomycin and ciprofloxacin, associated with simultaneous reduction of *Clostridiales* (in particular *Ruminococcaceae* and *Lachnospiraceae*), similar to what has previously been observed in adult patients [42]. Most interestingly, we found that treatment with vancomycin and ciprofloxacin was not only associated with reduced *Clostridiales*, but also with increased abundances of facultative anaerobic bacteria, e.g., *Enterobacteriaceae* (gamma-*Proteobacteria*). Vancomycin-treatment in a cohort of rheumatoid arthritis patients was previously shown to be associated with an expansion of *Proteobacteria* [43]. The use of a prophylactic ciprofloxacin treatment to prevent chemotherapy- and transplantation-related bloodstream infections is an established method [44], and it was proposed that fluoroquinolones (the antibiotic class comprising ciprofloxacin) can prevent intestinal domination of *Proteobacteria* in HSCT patients [15]. However, ciprofloxacin treatment in healthy subjects has been associated with decreased microbial diversity and decreased *Ruminococcaceae* and *Lachnospiraceae* abundances [45], indicating microbial community disruption. Therefore, our findings further challenge the choice of antibiotics, such as vancomycin and ciprofloxacin, in patients undergoing chemotherapy and HSCT. Interestingly, in the present study a number of less frequently used antibiotic agents, e.g. ceftazidime (a cephalosporin), showed positive associations with high *Clostridiales* abundances. In agreement, another cephalosporin (cefepime) has previously been attributed clostridial sparing effects [46]. However, ceftazidime was associated with reduced bacterial alpha diversity to a similar degree as vancomycin and ciprofloxacin in a previous study [42]. Elucidating the effects of antibiotic agents potentially contributing to maintaining gut microbial homeostasis in HSCT therefore require further investigation.Of note, we did not take the mode of application of the antibiotics into account here, which might limit the strength of our conclusion. It remains to be determined whether certain antibiotics modulate the patients’ microbiome and how this might lead to either positive or adverse clinical outcomes.

## Conclusions

Our findings support the increasing evidence of microbial involvement in the context of HSCT in cancer patients. We provide evidence for the association between specific microbial taxa and host immune markers. In particular, we examined the prognostic potential of immune markers and gut microbial community dynamics for immune reconstitution and outcomes after HSCT by revealing multivariate associations. We observed increased human beta-defensin 2 in patients with moderate to severe aGvHD and high mortality. In those patients, NK and B cell reconstitution was slow compared to patients with low mortality. These associations only applied when distinct gut microbial abundance patterns were observed, namely low abundances of *Ruminococcaceae* and high abundances of *Lactobacillaceae*. Therefore, hBD2, in connection with longitudinal microbial community pattern surveillance, could be further evaluated as a potential novel candidate marker to identify patients at risk of adverse outcomes (e.g. aGvHD) and slow immune cell reconstitution after HSCT, contributing to improved clinical outcomes. Of note, our cohort comprised a relatively small number of patients with different primary diseases and conditioning regimens, and our findings would therefore benefit from being assessed in larger, more homogenous patient groups. Importantly, microbial abundances also depended on antibiotic treatment. Our findings suggest that certain antimicrobial agents might contribute to a shift from obligate to facultative anaerobes. This highlights the need to assess the usage of specific antibiotics in more detail and to take antibiotic treatment into consideration when describing microbial communities in HSCT recipients.

## Methods

### Patient recruitment and sample collection

We recruited 37 children (age range 1.1 – 18.0 years) undergoing their first myeloablative allogeneic hematopoietic stem cell transplantation at Copenhagen University Hospital Rigshospitalet, Denmark, from June 2010 to September 2012. Patients’ clinical characteristics are listed in Additional file 1: Table S1. Further information can also be found in previous studies where the cohort has been examined in relation to other questions [17, 47–49]. All patients received pretreatment with a myeloablative conditioning regimen, starting on day-7 (Additional file 1: Table S1). Four patients were re-transplanted at day +157, +518, +712 and +1360 after the first transplantation, respectively.

Sampling time points were defined according to the following intervals: pre-HSCT (collected between day −33 and day −3), at the time of HSCT, preferably before graft infusion (collected between day −2 day +2) and weekly during the first six weeks after transplantation (week +1: day +3 to day +10, week +2: day +11 to day +17, week +3: day +18 to day +24, week +4: day +25 to day +31, week +5: day +32 to day +38, week +6: day +39 to day +45) (Figure 1A). Broader intervals applied to follow-up time points: Month +1 (between days +21 and +44), month +2 (between days +45 and +70), month +3 (between days +77 and +105), month +6 (between days +161 and +197) and 1 year post transplantation (between days +346 and +375).

### Infections and antibiotics

Bacterial infections from before transplantation until 1 year post-HSCT were taken into consideration for downstream analysis. For each time point, it was recorded whether any bacterial infection occurred within the respective specified interval or not (1/0). Antibiotic treatment from before HSCT (from day −90) until month +2 was taken into consideration. We included only those time points corresponding to the time points of microbiota profiling into downstream analyses.

### Analysis of T, B and NK cells in peripheral blood

T, B and NK cell counts were determined in months +1, +2, +3, and +6 post transplantation. Lymphocyte subsets in peripheral blood were quantified using Trucount Tubes (Becton Dickinson, Albertslund, Denmark) together with the following panel of conjugated monoclonal antibodies and analysed on a FC500 flow cytometer (Beckman Coulter, Copenhagen, Denmark): CD3-PerCP, CD3-FITC, CD4-FITC, CD8-PE, CD45-PerCP, CD16/56-PE, CD20-FITC and CD19-PE (Becton Dickinson). CD3+ T cells, CD3+CD4+ T cells and CD3+CD8+ T cells were determined. NK cells were differentiated by CD3-CD45+CD16+CD56+ phenotype. The following B cell phenotypes were distinguished: total B cells (CD45+CD19+), mature B cells (CD45+CD19+CD20+) and immature B cells (CD45+CD19+CD20−). Data of these immune cell populations have been published previously in a different context [47].

### Analysis of monocytes and neutrophils

Leukocyte numbers and subsets were monitored daily during hospitalization and subsequently every week in the outpatient clinic using flow cytometry (Sysmex XN) or, in case of very low leucocytes, counted by microscopy (CellaVision DM96 microscope). Mean monocyte and neutrophil counts were calculated for further analysis per time point according to the intervals specified above.

### Quantification of inflammatory markers

EDTA-anticoagulated and heparinized blood was sampled and then centrifuged within 2 hours after collection. The plasma was isolated and cryopreserved in 0.5-ml aliquots at −80°C. IL-6 levels on day +7 were determined in EDTA-anticoagulated plasma using the Human Th1/Th2/Th17 Cytometric Bead Array kit (Becton, Dickinson and Co., Denmark) and a FACSCalibur flowcytometer (Becton, Dickinson and Co), according to the manufacturer’s instructions with a detection limit of 2.5 pg/mL. IL-6 data have been published previously in another context for a larger cohort than the patients included here [17, 48]. CRP levels were measured daily by Modular P Modular (normal range, 0 to 10 mg/L) at the Department of Clinical Biochemistry, Copenhagen University Hospital Rigshospitalet, Denmark. Mean CRP levels were calculated for further analysis per time point according to the intervals specified above. Mean CRP levels pre-HSCT include measurements from day −7 to day −3, i.e. the days after the start of conditioning (except for day −7).

### Quantification of Citrulline

Plasma citrulline concentrations pre-HSCT (before the start of conditioning (day −7)) and at days +7 and +21 were measured by reverse-phase high-performance liquid chromatography of their phenylisothiocyanate derivatives from heparinized plasma, as described previously [17, 48]. Citrulline levels have previously been described for patients of this cohort in a different context [17, 48].

### Enzyme-linked immunosorbent assay (ELISA) of human beta defensins

Human beta defensin 2 and 3 (hBD2 and hBD3) concentrations in heparinized plasma samples of 37 patients at eight time points (pre-HSCT (before start of conditioning, except for 4 patients sampled at day −6 or −5), on the day of transplantation, weeks +1 to +4, month +2 and +3) and 10 healthy controls (sampled once each) were quantified by two-step sandwich ELISA following the manufacturer’s instructions (Peprotech Human BD-2 and BD-3, Standard ABTS ELISA Development Kit, cat.no. 900-K172 and 900-K210, respectively). Samples of 3 out of 10 healthy individuals were additionally spiked to a peptide concentration of 1000 pg/ml. Two replicates were measured per sample and their mean was used for further analysis. Samples were measured undiluted as well as in 1:4 and 1:16 dilutions to also cover concentrations potentially exceeding the upper detection limit. Samples with very high concentrations were additionally measured in 1:32 and 1:128 dilutions. Detection limits were 16 – 2000 pg/ml for hBD2 and 31– 4000 pg/ml for hBD3. Absorbance was measured on a VICTOR™ X3 Multilabel Plate Reader (Perkin Elmer, Inc., USA) at 405 nm. Wavelength correction at 540 nm was used to prevent optical interference caused by the material of the microtiter plate. Concentrations of hBD3 were mostly below the limit of detection, except for a few exceptionally high measurements (average: 1450.69 pg/ml, median: 0 pg/ml, range: 0 – 279038.71 pg/ml).

### DNA isolation from fecal samples and 16S rRNA gene sequencing

Fecal samples for analysis of the intestinal microbiome were collected from a subset of 30 patients at 7 time points: pre-HSCT (5 patients were sampled after the start of conditioning (between day −6 and day −4)), at the time of HSCT and once weekly during the first five weeks after transplantation. The intestinal microbiome was characterized at 1-2 time points in 8 patients (27%), at 3-4 time points in 15 patients (50%) and at 5-6 time points in 8 patients (27%) (Additional file 1: Table S1). In patients who underwent re-transplantation, no feces samples collected after the second transplantation were included in this study. In total, 97 fecal samples were obtained.

DNA from fecal samples and a blank control were isolated with the use of the Maxwell 16 Instrument (Promega Corporation) following the manufacturer’s instructions for the low elution volume blood DNA system. Alterations to the protocol included additional lysozyme treatment and bead beating with stainless steel beads for 2 min/20 Hz in a tissue lyser (Qiagen). In each sample including the blank control, the V4-V5 region of the 16S ribosomal RNA gene were amplified in PCR using the following barcoded primers: 519F (5#-CAGCAGCCGCGGTAATAC-3#) and 926R (5#-CCGTCAATTCCTTTGAGTTT-3#). Amplicons were then analyzed for quantity and quality in an Agilent 2100 Bioanalyzer (Agilent Technologies) with the use of an Agilent RNA 1000 Nano Kit. For library preparation, 50 ng of DNA from each sample was pooled with multiplex identifiers for 2-region 454 sequencing on GS FLX Titanium PicoTiterPlates (70675) with the use of a GS FLX Titanium Sequencing Kit XLR70 (Roche Diagnostics). Library construction and 454 pyrosequencing were performed at the National High-Throughput DNA Sequencing Centre, University of Copenhagen.

### 16S rRNA gene sequence pre-processing

Raw 454 sequence reads stored in standard flowgram format (SFF) were extracted, converted to and stored in FASTA format with associated quality files (containing sequence quality scores) using the *sffinfo* command of the bioinformatics software tool mothur [50]. Trimming according to the clipQualLeft and clipQualRight values provided by the sequence provider was disabled because cut-off values are opaque and not customizable.

Analysis was continued in the Quantitative Insights Into Microbial Ecology (QIIME; version 1.9.0) bioinformatics pipeline [51]. FASTA files were demultiplexed and quality filtered using the script *split_libraries.py* (Mapping files are available from figshare (https://dx.doi.org/10.6084/m9.figshare.6508250). As the samples were sequenced bidirectional, each FASTA file was demultiplexed in two steps. Firstly, based on a mapping file containing the 519F primer as the “LinkerPrimerSequence” and the 926R primer as the “ReversePrimer”, both in 5’ to 3’ orientation. Secondly, based on a mapping file containing the 926R primer as the “LinkerPrimerSequence” and the 519F primer as the “ReversePrimer”, again both in 5’ to 3’ orientation.

Reads between 200 and 1000 bp length and a minimum quality score of 25 were retained (default). Sequences with homopolymers longer than 200 bp were removed from the data set. Removal of reverse primer sequences (*-z truncate_only* option) was disabled during demultiplexing. Subsequently, the demultiplexed FASTA files that were not yet primer-truncated were then used to denoise flowgrams (.sff.txt files also generated by mothur’s *sffinfo*) with QIIME’s *denoise_wrapper.py* script. Reverse primer-truncation had not been done yet to ensure compatibility between FASTA and .sff.txt files. The denoised FASTA output files were then inflated, i.e., flowgram similarity between cluster centroids was translated to sequence similarity, to be used for OTU picking. Reverse primers and subsequent sequences in the demultiplexed and denoised FASTA files were then truncated using the *truncate_reverse_primer.py* script. In the following step, the orientation of the primer-truncated reads that started with the 926R primer as the “LinkerPrimerSequence” was adjusted by reverse complementation (with the script *adjust_seq_orientation.py*). All trimmed reads were then concatenated to a single file for further analysis.

Chimeras were identified using the script *identify_chimeric_seqs.py* and method *usearch61*, which performs both de novo (abundance-based) and reference-based detection (by comparing the dataset to the chimera-free reference database Ribosomal Database Project (RDP; training database version 15)). Only those sequences that were flagged as non-chimeras from both detection methods were retained (option *–non_chimeras_retention = intersection*). Operational taxonomic unit (OTU) clustering was performed, using the script *pick_otus.py* (based on the SILVA database, *Silva_119_rep_set97)*. OTU tables in BIOM format were created with *make_otu_table.py* (and subsequently converted to JSON BIOM format to be compatible with analysis in R [52] with the package *phyloseq* [53]). The OTU table and the taxonomy table are available from figshare (https://dx.doi.org/10.6084/m9.figshare.6508187).

### Statistical analyses

Statistical analyses and creation of graphs were performed with the program R (Version 3.4.0, R Foundation for Statistical Computing, Vienna, Austria) [52]. All R scripts documenting our statistical analyses are available from figshare (https://dx.doi.org/10.6084/m9.figshare.6508238). Sequencing data and all related experimental and clinical data (data sets available from figshare, https://dx.doi.org/10.6084/m9.figshare.6508232) were integrated for analysis with the R package *phyloseq* and its dependencies [53] (Additional file 11). The resulting phyloseq objects are provided through figshare (https://dx.doi.org/10.6084/m9.figshare.6508235). Plots were generated with the packages *ggplot2* [54], *plotly* [55], and *mixOmics* [56]. Dose-response analysis of the ELISA data was performed with four-parameter log-logistic models in the R package *drc* [57].

Alpha diversity (measured by inverse Simpson index), levels of human beta-defensin 2 (hBD2) concentration, citrulline and C-reactive protein (CRP), as well as monocyte counts, NK cell counts, total B cell counts, and CD4+ T cell counts at different time points were compared using Friedman tests with Benjamini-Hochberg correction for multiple testing (Additional files 3: Figure S2, Additional files 11 and 12). In addition, Kendall’s coefficient of concordance (Kendall’s *W*) was calculated on ranked data for each marker (Additional file 13). Like the Friedman test, the test for Kendall’s *W* allowed the comparison of marker levels between time points. In addition, the coefficient of concordance informs about the level of agreement between patients. Therefore, Kendall’s *W* can be interpreted as a measure of effect size for the Friedman tests. A Kendall’s *W* <0.1 was considered as indicating a small effect, 0.1 – 0.5 as a moderate effect, and > 0.5 as a strong effect. As an exception, hBD2 in healthy controls was compared with hBD2 in patients at individual time points by using Wilcoxon rank-sum tests, because hBD2 was only measured once in the healthy control individuals and can therefore not be analyzed in a Friedman test designed for repeated measurements. Monocyte counts in Additional file 2: Figure S1B are depicted at more time points than indicated in Figure 1A because not all time points were included into further analyses. The day of HSCT, week +1 and week +2 were excluded as data were missing for ≥40% of the patients. Months +2, +3, and +6, as well as 1 year were chosen as representative follow-up time points for further analyses as indicated in Figure 1A.

A core set of OTUs was obtained by retaining 256 OTUs (out of 756 OTUs) with ≥5 reads in ≥ 2 samples using the function *kOverA()* from R package *genefilter* [58]. Subsequently, 17 OTUs that were more abundant in the blank control than in the majority of samples were removed as potential contaminants prior to downstream analyses. The resulting count data set of 239 OTUs was transformed for subsequent analyses using the function *varianceStabilizingTransformation()* in the package *DESeq2* [59] (Additional file 11). The function implements a Gamma-Poisson mixture model [60] to account for both library size differences and biological variability.

Median imputations were performed for continuous clinical and immune marker data with less than 20% missing values. Variables with more than 20% missing values were excluded from the analysis (Additional files 13 and 14). Central tendencies of immune cell counts at single time points in relation to clinical outcomes (maximum aGvHD grade 0-I vs. grade II-IV) were assessed in univariate analyses by Wilcoxon rank-sum tests and displayed in boxplots (Additional file 14).

A model selection procedure was implemented to find the relevant variables to be included in subsequent multivariate analyses of how microbiome patterns are associated with clinical outcomes, baseline parameters and immune parameters during the course of transplantation: A Manhattan distance matrix of the variance-stabilized bacterial community data was calculated using the *distance()* function in *phyloseq* [53]. Subsequently, permutational multivariate analysis of variance using distance matrices (adonis) for model selection was performed by applying the *adonis2()* function in the package *vegan* [61] (Additional files 3: Figure S2, Additional file 15). Permutation design was set up with respect to repeated measurements within the same patients and the intact chronological order of time points. Besides immune marker levels and immune cell counts (pre- and post-HSCT, i.e. recipient- and donor-derived cells, respectively) at the time points described above, we included clinical outcomes (i.e. overall survival, aGvHD (grade 0-I vs II-IV), and relapse) after transplantations, antibiotic treatment during the course of transplantation and clinical patient characteristics in the model to account for possible effects of recipient age at the time of transplantation, recipient sex, donor type (sibling vs. unrelated), malignant vs. benign diagnosis, graft type (stem cell source: bone marrow, umbilical cord blood or peripheral blood) and application of irradiation therapy (yes/no). Variables that were found to be significant (*P* ≤ 0.05) in the adonis analysis were included in subsequent multivariate multi-table analyses, i.e., sparse partial least squares (sPLS) regression and canonical correspondence analysis (CCpnA) (Additional file 15). Choosing variables with significant effects in adonis for follow-up statistical testing has been performed previously [62]. Even though validation of the set of selected variables through a data-splitting approach might be preferable, this was not feasible due to our relatively small data set. To account for this and to avoid post-selective inference, we renounce calculation of p-values from the two analyses that directly depend on the pre-selection (sPLS and CCpnA).

Correlations among the selected clinical variables were assessed in correlation matrices based on Spearman’s rank correlation tests (Additional file 3: Figure S2, Additional file 14). Matrices were calculated using the *rcorr()* function of the R package *Hmisc* [63] and displayed with the package *corrplot* [64]. *P*-values were calculated with the *rcorr.adjust()* function with correction for multiple testing (method “Holm”). The Spearman’s rank correlation tests were performed on the set of variables selected from the adonis analysis. However, here we assess correlations among those variables, and not between variables and microbial abundances.

Sparse PLS regression was performed by applying the *spls()* function in the package *mixOmics* [56] (Additional file 3: Figure S2, Additional file 15). The sPLS regression allows the integration of the microbial community data matrix and the clinical variable matrix for multiple regression. It is robust enough to handle collinearity and noise in the data and is suitable to model multiple response variables [65]. The number of clinical variables to be kept in the model for each component (*keepY*) was set to 23, corresponding to the number of variables pre-selected with adonis. We ran the sPLS regression with a range of numbers (20-40) of OTUs to be kept for each component (*keepX*). As the results were robust to this choice, *keepX* was set to 30. The number of components to choose was estimated with the *perf*() function and set to *ncomp* = 2. The sPLS model was run in regression mode. Thereafter hierarchical clustering was performed within the mixOmics *cim()* function based on the sPLS regression model with the clustering method “complete linkage” and the distance method “Pearson’s correlation”. Coefficients of pairwise correlations between OTU abundances and clinical variables were thereby obtained. Furthermore, loading plots were generated with the function *plotLoadings()* (method = “mean”) to visualize loading vectors of specific OTUs that contribute most to the separation of variables in components 1 and 2.

Canonical (i.e., bidirectional) correspondence analysis (CCpnA), a multivariate constrained ordination method, was performed by using the *cca()* function in the package *vegan* [61] (Additional file 3: Figure S2, Additional file 15). In this method, the microbial community data matrix is Chi-square transformed and weighted linear regression on pre-selected constraining variables is performed. The resulting fitted values are used for correspondence analysis by singular value decomposition. CCpnA is a constrained method in the sense that it does not aim at depicting all variation in the data, but only the variation directly explained by the constraints (i.e., the provided set of pre-selected variables). The resulting triplot is not displayed as a square representation, but rather corresponds to the percentage of variance explained by axis 1 and 2, respectively, as previously suggested [66]. In contrast to the sPLS analysis, the CCpnA was performed in canonical mode, i.e., modeling bidirectional relations between OTU abundances and clinical variables. OTUs with a correlation of >0.2/<−0.2 in the sPLS analysis were included in the CCpnA model.

As another approach to distinguish between microbial community states of the intestinal microbiome, we assigned samples to community state types (CSTs) by partitioning around medoid (PAM) clustering (function *pam()* in package *cluster* [67]) based on a Jensen-Shannon distance of the variance stabilized microbial count data (R code modified after [68]) (Additional file 3: Figure S2, Additional file 16). The number of clusters was determined by gap statistic evaluation and silhouette width quality validation. We further assessed patients’ transitions between CSTs over time. OTUs were assigned to CST - based clusters (Additional file 4: Table S2) based on in which CST they exhibited the highest average abundance over all samples (within each CST). Furthermore, we showed detailed longitudinal profiles of the microbial community on family-level, and selected immune markers for individual patients with aGvHD (Additional file 17).

## List of abbreviations

3-IS: 3-indoxyl sulfate
aGvHD: Acute graft-versus-host disease
ALL: Acute lymphoblastic leukemia
AML: Acute myeloid leukemia
AMP: Antimicrobial peptide
CCpnA: Canonical correspondence analysis
CRP: C-reactive protein
CST: Community state type
ELISA: Enzyme-linked immunosorbent assay
GvL effect: Graft-versus-leukemia effect
hBD2/hBD3: Human beta-defensin 2/3
HLA: Human leukocyte antigen
HSCT: Hematopoietic stem cell transplantation
IL-6: Interleukin-6
NK cell: Natural killer cell
OTU: Operational taxonomic unit
P: Patient
PAM clustering: Partitioning around medoid clustering
QIIME: Quantitative Insights Into Microbial Ecology
SFF: Standard flowgram format
sPLS regression: sparse partial least squares regression
TBI: Total body irradiation

## Declarations

## Acknowledgements

We thank the patients and their families for participating in the study, and the National High-Throughput DNA Sequencing Centre at the University of Copenhagen for library construction and sequencing. Furthermore, we are grateful for helpful discussions about the statistical analyses with Pratheepa Jeganathan and Kris Sankaran from the Department of Statistics at Stanford University. We thank Jeffrey Edward Skiby for reviewing the manuscript.

## Availability of data and material

The 16S rRNA gene sequences are available through the European Nucleotide Archive (ENA) at the European Bioinformatics Institute (EBI) under accession number PRJEB25221. The datasets generated and/or analysed during the current study as well as the R code used to analyze the data are available from the figshare repository at https://figshare.com/projects/Specific_gut_microbiome_members_are_associated_with_distinct_immune_markers_in_allogeneic_hematopoietic_stem_cell_transplantation/35201.

## Author’s contributions

A.C.M., K.K., K.G.M., and S.J.P. designed the research; A.C.M., K.K., M.S.C., and S.J.P. performed the research; A.C.M., S.H., and S.J.P. contributed analytic tools; A.C.M., and S.J.P. analysed the data; A.C.M. and S.J.P. wrote the manuscript; and K.K., S.H., F.M.A., O.L., and K.G.M. edited the manuscript. All authors have read and approved the manuscript as submitted.

## Ethics approval and consent to participate

Written informed consent was obtained from the patients and/or their legal guardians. The study protocol was approved by the local ethics committee (H-1-2010-009) and the Danish Data Protection Agency.

## Consent for publication

Not applicable.

## Additional files

**Additional file 1: Table S1.** Clinical patient characteristics. General patient characteristics, conditioning regimens, complications, and outcomes for the pediatric cohort (*n=37*) and the subcohort (*n=30*) for which the intestinal microbiome was characterized. Abbreviations: HLA, human leukocyte antigen; TBI, total body irradiation; CY, Cyclophosphamide; VP16, Etoposide; BU, Busulfan; MEL, Melphalan; GvHD, graft-versus-host disease. (PDF)

| Characteristics |  | Number of patients (all) | Percentage of all patients (%) | Number of patients (Sub-cohort for which the microbiota was characterized) | Percentage of sub-cohort (%) |
| --- | --- | --- | --- | --- | --- |
| Transplant recipients |  | 37 | 100 | 30* | 81.1 |
| Average recipient age in years |  | 8.2 (Range: 1.1-18.0) | NA | 7.8 (Range: 1.1-16.5) | NA |
| Intestinal microbiome characterized | At 1-2 timepoints | NA | NA | 8 | 26.7 |
|  | At 3-4 timepoints | NA | NA | 15 | 50 |
|  | At 5-6 timepoints | NA | NA | 8 | 26.7 |
| Patient sex | Female | 15 | 40.5 | 12 | 40 |
|  | Male | 22 | 59.5 | 18 | 60 |
| Disease at transplantation | malignant hematologic diseases | 23 | 62.2 | 17 | 56.7 |
|  | Severe aplastic anemia | 5 | 13.5 | 5 | 16.7 |
|  | Other benign disorders including immunodeficiencies | 9 | 24.3 | 8 | 26.7 |
| Donor type | HLA-matched sibling | 7 | 18.9 | 6 | 20 |
|  | HLA-matched unrelated donor (9/10 or 10/10 match) | 27 | 73 | 22 | 73.3 |
|  | HLA-mismatched umbilical cord blood donor (8/10 match) | 3 | 8.1 | 2 | 6.7 |
| Stem cell source | Bone marrow | 30 | 81.1 | 25 | 83.3 |
|  | Umbilical cord blood | 4 | 10.8 | 3 | 10 |
|  | Peripheral blood | 2 | 5.4 | 1 | 3.3 |
|  | Bone marrow and umbilical cord blood | 1 | 2.7 | 1 | 3.3 |
| Conditioning regimen | TBI + CY / TBI + VP16 | 10 | 27.1 | 10 | 33.3 |
|  | Combinations of BU, CY, VP16 and MEL | 18 | 48.7 | 12 | 40 |
|  | Fludarabine-based conditioning | 9 | 24.3 | 8 | 26.7 |
| **Anti-thymocyte globulin treatment** |  | 28 | 75.7 | 23 | 76.7 |
| **Antibiotics pre- and post-HSCT** |  | 37 | 100 | 30 | 100 |
| **Sex mismatch (female donor to male recipient)** |  | 4 | 10 | 3 | 10 |
| **Acute GvHD** | **Grade 0-I** | 26 | 70.3 | 22 | 73.3 |
|  | **Grade II-IV** | 11 | 29.2 | 8 | 26.7 |
| **Chronic GvHD within 24 months** |  | 8 | 18.9 | 5 | 16.7 |
| **Bacterial infections** | **At least one registered bacterial infection** | 25 | 67.6 | 19 | 63.3 |
|  | **No registered bacterial infection** | 12 | 32.4 | 11 | 36.7 |
| **Overall Survival\*\*** | **Alive** | 30 | 81.1 | 24 | 80 |
|  | **Dead** | 7 | 18.9 | 6 | 20 |
| **Relapse of primary malignancy** |  | 6 | 16.2 | 7 | 23.3 |
| **Non-relapse mortality** |  | 2 | 5.4 | 1 | 3.3 |
| **Re-transplantation** |  | 3 | 8.1 | 4 | 13.3 |

\*29/37 +1, one patient's microbiome was characterized but for which hBD2 was not measured
\*\* Mean follow-up time after HSCT: 62.1 months (range: 48.9 – 75.8 months)

**Additional file 2: Figure S1.**
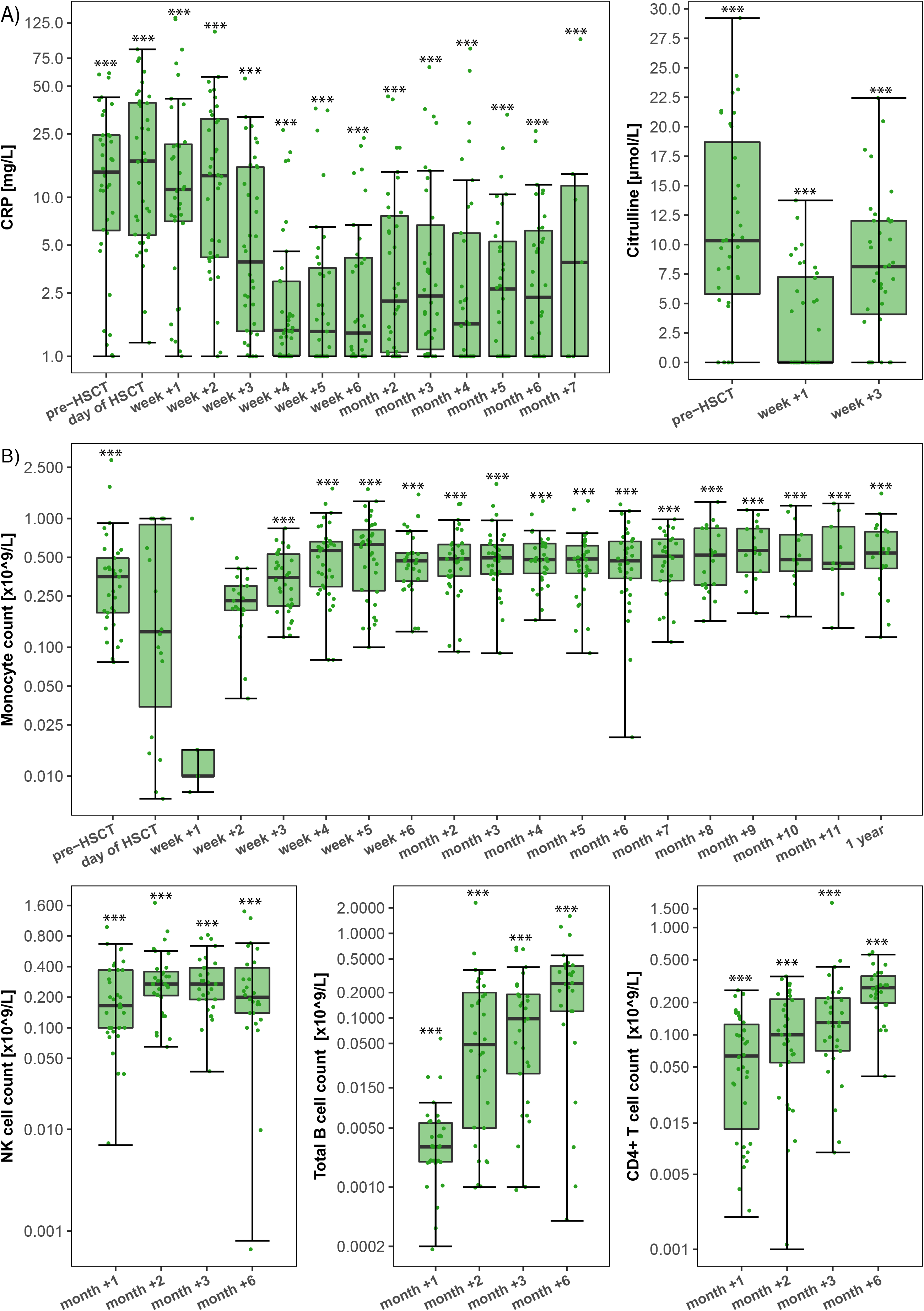
Temporal patterns of immune markers and immune cells in HSCT patients. (A) C-reactive protein (CRP) and plasma citrulline levels in HSCT patients (*n = 37*) over time. CRP levels were significantly higher prior to HSCT and until week +2 compared to all following time points, e.g. at the day of HSCT (median: 16.93 mg/L, range: 1.22 – 85.28 mg/L) compared to week +3 (median: 3.92 mg/L, range: 1.22 – 55.89 mg/L) (*P* < 0.001). Plasma citrulline levels were significantly lower in week +1 compared to pre-HSCT (*P* < 0.001) and week +3 (*P* < 0.001). (B) Immune cell counts in HSCT patients over time. Monocyte counts are depicted at more time points than indicated in Figure 1A, because not all time points were included into further analyses (see Methods). NK cell counts were higher in months +2 to +6 compared to in month +1 (*P* < 0.001). B cell counts as well as CD4+ T cell counts increased steadily from month +1 to month +6 (*P* < 0.001). Y-axes in all plots, except for citrulline, were log10-transformed for better visualization. Zeros were replaced with 1 to avoid undefined values on the log-transformed axes. Asterisks indicate whether the component at each respective time point was significantly different from any of the other time points (showing the maximum significance level). * *P* < 0.05, ** *P* < 0.01 and *** *P* < 0.001. (PDF)

**Additional file 3: Figure S2.**
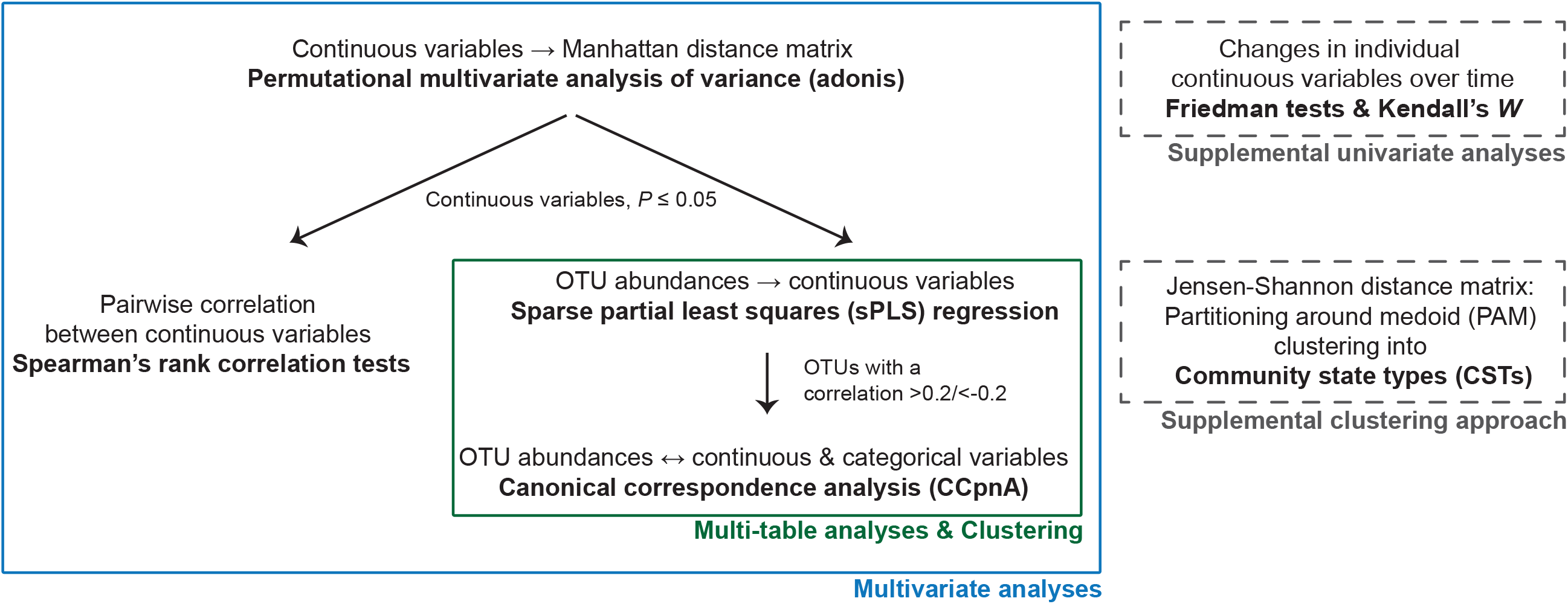
Workflow of the statistical analysis approach. The diagram displays the major steps of the statistical analyses and their dependencies. Multivariate analyses (blue box) constitute the main approach, especially the multi-table analyses and clustering analyses (green box). To unravel the complexity of the multivariate analyses, these were supplemented with univariate analyses (upper grey box). (PDF)

**Additional file 4: Table S2.** Results of Permutational Multivariate Analysis of Variance Using Distance Matrices (adonis). Adonis was employed for model selection to identify relevant immune markers and immune cell types to be included in downstream analyses (See Methods for details). Significant variables (P<0.05) are marked in bold. Abbreviations: hBD2_sim, plasma human beta-defensin 2 levels at time points simultaneous to microbiome characterization; CRP_sim, C-reactive protein levels at time points simultaneous to microbiome characterization; Lymphocyte_count_sim, total lymphocyte counts at time points simultaneous to microbiome characterization; pIL6, plasma interleukin 6 concentration; Citr, plasma citrulline concentration; CD3+, CD3+ T cell counts; CD4+, CD3+CD4+ T cell counts; CD8+, CD3+CD8+ T cell counts; B, total B cell (CD45+CD19+) counts; mat_B, mature B cell (CD45+CD19+CD20+) counts. immat_B, immature B cell (CD45+CD19+CD20−) counts; NK, natural killer cell counts; mean_mono, mean monocyte counts at indicated time point; mean_neutro, mean neutrophil counts at indicated time point; Timepoints: pre, prior to transplantation; w0, on the day of transplantation; w1, w2, w3, w4, w5: one, two, three, four and five weeks after transplantation, respectively; m1, m2, m3, m4, m6: one, two, three, four and six months after transplantation, respectively; 1y, 1 year post transplantation. (PDF)

| Variable | SumOfSqs | F | P-value |
| --- | --- | --- | --- |
| hBD2_sim | 9763 | 0.4635 | 0.915 |
| **hbd2_pre** | 89958 | 4.3373 | **0.001** |
| **hbd2_w0** | 39432 | 1.9012 | **0.037** |
| **hbd2_w1** | 62425 | 3.0098 | **0.007** |
| **hbd2_w2** | 70010 | 3.3755 | **0.004** |
| hbd2_w3 | 32106 | 1.5480 | 0.088 |
| hbd2_m2 | 33806 | 1.6300 | 0.104 |
| **CRP_sim** | 28707 | 1.3629 | **0.015** |
| CRP_pre | 15714 | 0.8102 | 0.631 |
| CRP_w0 | 22573 | 1.1639 | 0.280 |
| **CRP_w1** | 51055 | 2.6324 | **0.011** |
| CRP_w2 | 34122 | 1.7593 | 0.098 |
| CRP_w3 | 41269 | 2.1278 | 0.055 |
| CRP_m1 | 27106 | 1.3976 | 0.169 |
| **CRP_w5** | 41965 | 2.1637 | **0.026** |
| **CRP_w6** | 38860 | 2.0036 | **0.029** |
| CRP_m2 | 31527 | 1.6255 | 0.095 |
| **CRP_m3** | 39972 | 2.0610 | **0.034** |
| CRP_m4 | 23961 | 1.2354 | 0.252 |
| **CRP_m6** | 39945 | 2.0596 | **0.024** |
| plL6_w1 | 23180 | 1.1431 | 0.311 |
| **Citr_pre** | 78105 | 3.8519 | **0.002** |
| **Citr_w1** | 42029 | 2.0727 | **0.021** |
| Citr_w3 | 33872 | 1.6705 | 0.069 |
| CD3+_m1 | 28654 | 1.4500 | 0.123 |
| CD3+_m2 | 17891 | 0.9054 | 0.501 |
| CD4+_m1 | 30266 | 1.5316 | 0.090 |
| **CD4+_m2** | 38132 | 1.9296 | **0.037** |
| **CD8+_m1** | 38615 | 1.9541 | **0.029** |
| CD8+_m2 | 21275 | 1.0766 | 0.337 |
| B_m1 | 26588 | 1.3454 | 0.166 |
| **B_m2** | 41803 | 2.1154 | **0.025** |
| mat_B_m1 | 26622 | 1.3472 | 0.167 |
| **mat_B_m2** | 41697 | 2.1100 | **0.027** |
| immat_B_m1 | 28078 | 1.4208 | 0.133 |
| **immat_B_m2** | 40142 | 2.0313 | **0.029** |
| **NK_m1** | 63891 | 3.2331 | **0.002** |
| **NK_m2** | 44566 | 2.2552 | **0.020** |
| Lymphocyte_count_sim | 21541 | 1.0227 | 0.610 |
| Lymphocyte_m2 | 22196 | 1.1232 | 0.324 |
| Lymphocyte_m3 | 24450 | 1.2373 | 0.224 |
| Lymphocyte_m6 | 26278 | 1.3298 | 0.177 |
| Lymphocyte_1y | 29658 | 1.5008 | 0.114 |
| **mean_mono_pre** | 47349 | 2.3109 | **0.013** |
| **mean_mono_w3** | 64242 | 3.1353 | **0.002** |
| mean_mono_m1 | 17924 | 0.8748 | 0.561 |
| mean_mono_m2 | 25930 | 1.2655 | 0.187 |
| mean_mono_m3 | 18269 | 0.8916 | 0.516 |
| mean_mono_m6 | 1579 | 0.7708 | 0.654 |
| mean_neutro_pre | 25198 | 1.2298 | 0.225 |
| mean_neutro_w3 | 31161 | 1.5208 | 0.149 |
| mean_neutro_m1 | 29036 | 1.4171 | 0.137 |
| mean_neutro_m2 | 16656 | 0.8129 | 0.611 |
| **mean_neutro_m3** | 48466 | 2.3653 | **0.010** |
| mean_neutro_m6 | 15960 | 0.7789 | 0.653 |

**Additional file 5: Table S3:** Taxonomy and cluster affiliation of OTUs strongly associated with host-related variables based on sPLS analysis and community state typing (CST). List of the 57 OTUs correlated strongest with variables in the sPLS analysis (>0.2/<−0.2). SPLS-based clusters were determined by applying the mixOmics *cim()* function to the sPLS regression model (hierarchical clustering method: complete linkage, distance method: Pearson’s correlation) (see Methods). Four community state types (CSTs) were defined by clustering of fecal samples with similar microbial community compositions by partitioning around medoid (PAM) clustering (see Methods). OTUs were then assigned to the CST-based clusters in which they exhibited the highest average abundance over all samples. The same taxonomic families dominated in sPLS- and CST-based clusters, respectively. Cluster 1 was dominated by *Lactobacillaceae*. Cluster 2 was characterized mainly by *Ruminococcaceae* and *Lachnospiraceae*. Cluster 3 harbored Proteobacteria (P), e. g. *Enterobacteriaceae*. CS-typing revealed one additional cluster (4), characterized by a high abundance of *Enterococcaceae* and *Staphylococcaceae*. OTU numbers refer to the SILVA database (*silva_119_rep_set97)*. Phyla abbreviations: F, Firmicutes; B, Bacteroidetes; A, Actinobacteria; P, Proteobacteria; FU, Fusobacteria. (PDF)

| OTU number | Phylum | Order | Family | Genus/Species/Description | Cluster |  |
| --- | --- | --- | --- | --- | --- | --- |
|  |  |  |  |  | sPLS | CST |
| HM534767.1 | F | Lactobacillales | Lactobacillaceae | Lactobacillus sp. Akpobro1 | 1 | 1 |
| AF413523.1 | F | Lactobacillales | Lactobacillaceae | Lactobacillus pantheris | 1 | 1 |
| KF751750.1 | F | Lactobacillales | Lactobacillaceae | Lactobacillus sp. | 1 | 1 |
| AB257866.1 | F | Lactobacillales | Lactobacillaceae | Lactobacillus suebicus | 1 | 1 |
| JX986976.1 | F | Lactobacillales | Lactobacillaceae | Lactobacillus aviarius subsp. araffinosus | 1 | 1 |
| GU451064.1 | F | Lactobacillales | Lactobacillaceae | Lactobacillus sp. 5-1-2 | 1 | 1 |
| EU774037.1 | F | Lactobacillales | Lactobacillaceae | Lactobacillus sp. | 1 | 1 |
| FN667075.1 | F | Lactobacillales | Lactobacillaceae | Lactobacillus sp. | 1 | 1 |
| EU461951.1 | F | Lactobacillales | Streptococcaceae | Streptococcus sp. | 1 | 1 |
| EU460769.1 | F | Lactobacillales | Lactobacillaceae | Lactobacillus sp. | 1 | 1 |
| KC836559.1 | F | Lactobacillales | Streptococcaceae | Streptococcus salivarius subsp. thermophilus | 1 | 1 |
| FJ749794.1 | F | Lactobacillales | Lactobacillaceae | Lactobacillus fermentum | 1 | 1 |
| KF029502.1 | F | Lactobacillales | Lactobacillaceae | Lactobacillus casei | 1 | 1 |
| AUFL01000034 | B | Bacteroidales | Porphyromonadaceae | Dysgonomonas capnocytophagoides DSM 22835 | 1 | 1 |
| DQ801646.1 | F | Clostridiales | Lachnospiraceae | uncultured bacterium | 2 | 2 |
| EU246837.1 | F | Bacillales | Staphylococcaceae | Staphylococcus sp. Cobs2Tis23 | 2 | 4 |
| DQ804549.1 | F | Clostridiales | Ruminococcaceae | Faecalibacterium sp. | 2 | 2 |
| FJ366680.1 | A | Bifidobacteriales | Bifidobacteriaceae | Bifidobacterium sp. | 2 | 2 |
| HQ749687.1 | F | *Clostridiales* | *Ruminococcaceae* | uncultured bacterium | 2 | 2 |
| FJ753793.1 | F | *Clostridiales* | *Peptostreptococcaceae* | swine fecal bacterium RF1A-Xyl2 | 2 | 2 |
| DQ800353 | F | *Clostridiales* | *Lachnospiraceae* | *Blautia* sp. | 2 | 2 |
| EU887971.1 | F | *Clostridiales* | *Ruminococcaceae* | uncultured Clostridia bacterium | 2 | 2 |
| JQ608258.1 | F | *Erysipelotrichales* | *Erysipelotrichaceae* | bacterium NLAE-zl-C558 | 2 | 2 |
| AY850535.1 |  | *Bifidobacteriales* | *Bifidobacteriaceae* | *Bifidobacterium* sp. | 2 | 2 |
| DQ326608.1 | F | *Clostridiales* | *Ruminococcaceae* | uncultured bacterium | 2 | 2 |
| AJ272200.1 | F | *Lactobacillales* | *Enterococcaceae* | *Enterococcus hirae* | 2 | 4 |
| EU508251.1 | F | *Erysipelotrichales* | *Erysipelotrichaceae* | uncultured bacterium | 2 | 2 |
| DQ807523.1 | F | *Clostridiales* | *Lachnospiraceae* | uncultured bacterium | 2 | 2 |
| CP001726 | A | *Coriobacteriales* | *Coriobacteriaceae* | *Eggerthella lenta* DSM 2243 | 2 | 2 |
| HQ794871.1 | F | *Erysipelotrichales* | *Erysipelotrichaceae* | uncultured organism | 2 | 2 |
| JF193283.1 | F | *Clostridiales* | *Family XI* | *Finegoldia* sp. | 2 | 2 |
| DQ802363.1 | F | *Clostridiales* | *Lachnospiraceae* | *Blautia* sp. | 2 | 2 |
| GQ175428.1 | F | *Clostridiales* | *Ruminococcaceae* | uncultured bacterium | 2 | 2 |
| EU778851.1 | F | *Clostridiales* | *Ruminococcaceae* | *Subdoligranulum* sp. | 2 | 2 |
| GQ133038.1 | F | *Lactobacillales* | *Enterococcaceae* | *Enterococcus* sp. | 2 | 4 |
| JQ448431.1 | F | *Lactobacillales* | *Streptococcaceae* | *Streptococcus* sp. | 3 | 1 |
| JQ460192.1 | B | *Bacteroidales* | *Prevotellaceae* | *Prevotella* sp. | 3 | 3 |
| M58833.1 | F | *Lactobacillales* | *Lactobacillaceae* | *Pediococcus acidilactici* | 3 | 1 |
| HG799951.1 | F | Bacillales | Staphylococcaceae | Staphylococcus epidermidis | 3 | 3 |
| JF235878.1 | A | Micrococcales | Microbacteriaceae | Naasia sp. | 3 | 3 |
| JQ680457.1 | A | Corynebacteriales | Corynebacteriaceae | Corynebacterium sp. DYS15 | 3 | 3 |
| GQ379595.1 | P | Xanthomonadales | Xanthomonadaceae | Stenotrophomonas sp. | 3 | 4 |
| AB787271.1 | B | Bacteroidales | Bacteroidaceae | Bacteroides sp. UasXn-3 | 3 | 3 |
| AOTI010230528.6 | B | Flavobacteriales | Flavobacteriaceae | Chryseobacterium Triticum urartu | 3 | 3 |
| JF109069.1 | F | Bacillales | Staphylococcaceae | Staphylococcus sp. | 3 | 3 |
| JX464212.1 | F | Bacillales | Paenibacillaceae | Paenibacillus sp. BJC15-C21 | 3 | 3 |
| HQ778412.1 | B | Bacteroidales | Bacteroidaceae | Bacteroides sp. | 3 | 2 |
| FJ976590.1 | P | Enterobacteriales | Enterobacteriaceae | Enterobacter sp. LCR81 | 3 | 3 |
| KF178309.1 | F | Lactobacillales | Lactobacillaceae | Pediococcus pentosaceus | 3 | 3 |
| HQ703879.1 | F | Lactobacillales | Carnobacteriaceae | Alkalibacterium sp. | 3 | 3 |
| FJ950694.1 | P | Enterobacteriales | Enterobacteriaceae | Escherichia coli | 3 | 3 |
| JF164165.1 | F | Bacillales | Staphylococcaceae | Staphylococcus sp. | 3 | 3 |
| KC463799.1 | P | Enterobacteriales | Enterobacteriaceae | Klebsiella sp. | 3 | 3 |
| JF146381.1 | F | Lactobacillales | Streptococcaceae | Streptococcus sp. | 3 | 3 |
| AM900778.1 | F | Bacillales | Paenibacillaceae | Paenibacillus sp. PA215 | 3 | 3 |
| CBVB010000006 | B | Bacteroidales | Bacteroidaceae | bacterium MS4 | 3 | 3 |
| AB013258.1 | P | Neisseriales | Neisseriaceae | uncultured beta proteobacterium | 3 | 3 |

**Additional file 6: Figure S3.**
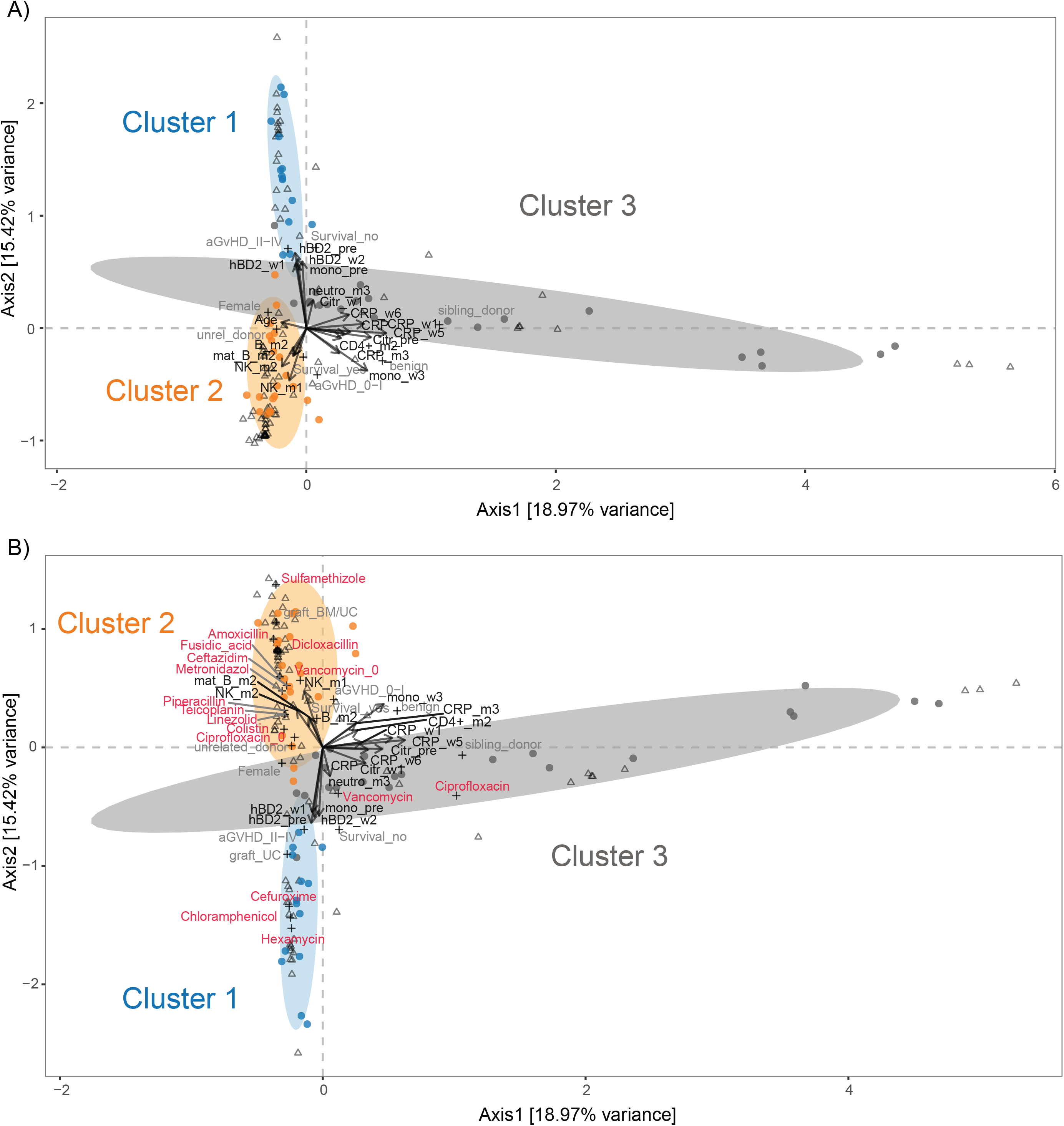
Canonical correspondence analysis (CCpnA) of immune markers and intestinal bacterial taxa in patients undergoing HSCT. Triplots showing dimension 1 and 2 of the CCpnA that includes continuous clinical variables (arrows), categorical variables (+), and OTUs (circles). Samples are depicted as triangles. OTUs with a correlation of >0.2/<−0.2 in the sPLS analysis were included in the CCpnA model. Only the variables and OTUs with a score >0.2/<−0.2 in at least one CCpnA dimension are shown. The OTUs in the CCpnA plots are colored according to the cluster they were affiliated with in the sPLS-based hierarchical clustering analysis, and the ellipses present an 80% confidence interval, assuming normal distribution. (A) Full size visualization corresponding to the CCpnA model shown in Figure 4. Plot dimensions correspond to the explained variances of each component. (B) CCpnA including antibiotic treatment at time points simultaneous to microbiome characterization. Antibiotics were added as categorical variables. Depiction of the antibiotic’s name (in red) indicates administration of the particular antibiotic, and the extension “_0” indicates no administration of the respective antibiotic. Abbreviations of variables are the same as in Figure 2. Further abbreviations: graft_BM: stem cell source bone marrow; graft_UC: stem cell source umbilical cord blood. (PDF)

**Additional file 7: Figure S4.**
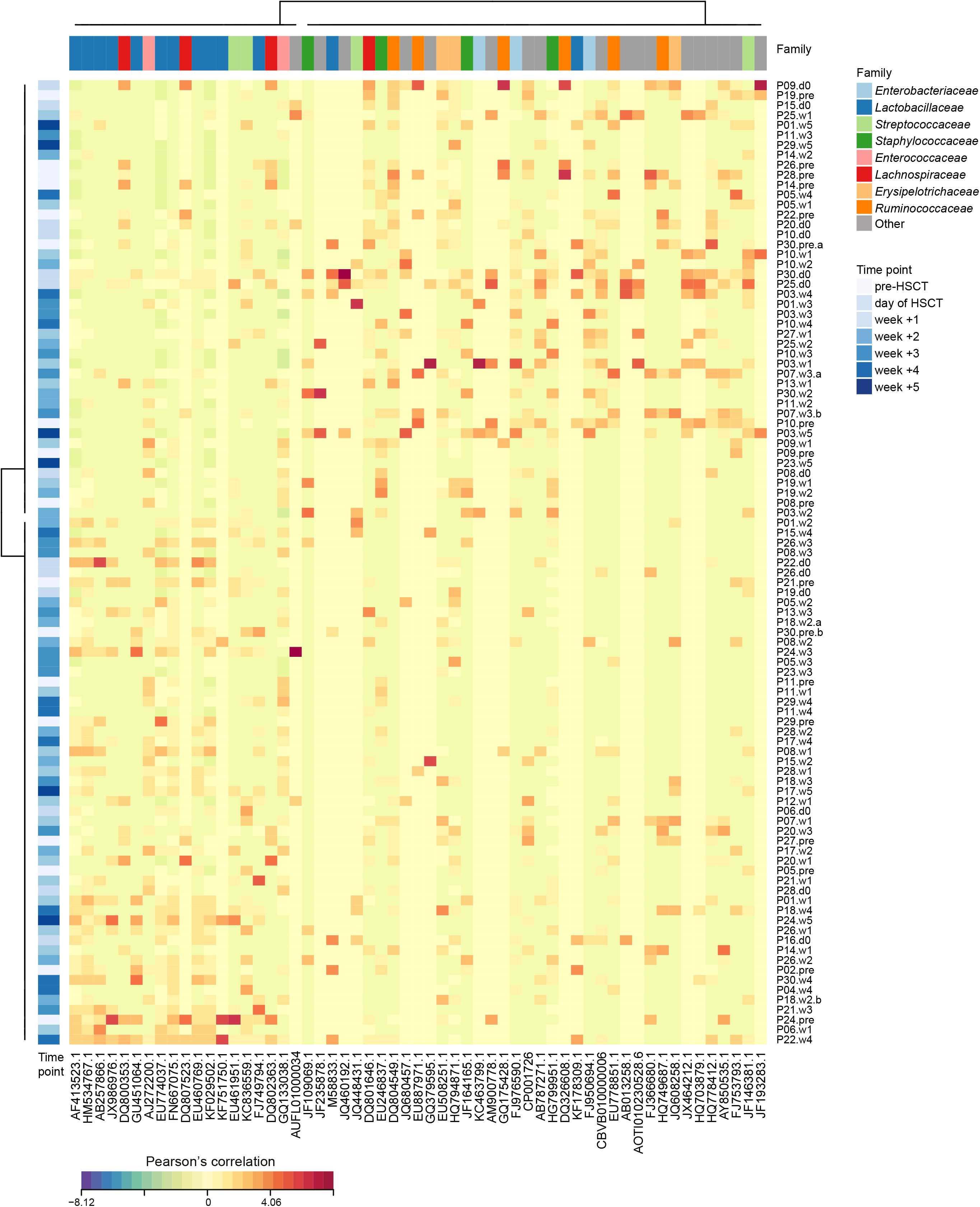
Clustered image map (CIM) of OTU abundances by patient in the first two sPLS dimensions. Hierarchical clustering of OTU abundances (bottom) and patients’ fecal samples (right) (clustering method: complete linkage, distance method: Pearson’s correlation) was performed within the mixOmics *cim()* function based on the sPLS regression model. High abundance of an OTU in a sample is represented as positive correlation in the map (red) and low abundance as negative correlation (blue). The sampling time points of the fecal samples are displayed in the side bar on the left (blue gradient from pre-HSCT time point (light blue) to week +5 post HSCT (dark blue)). The top side bar shows taxonomic information on family level. Sample names on the right indicate patient (P) and time point (pre: pre-HSCT, d0: day of HSCT, w: week). An “a” or “b” indicates that two samples were collected from the respective patient at the same time point, but on two different days. (PDF)

**Additional file 8: Figure S5.**
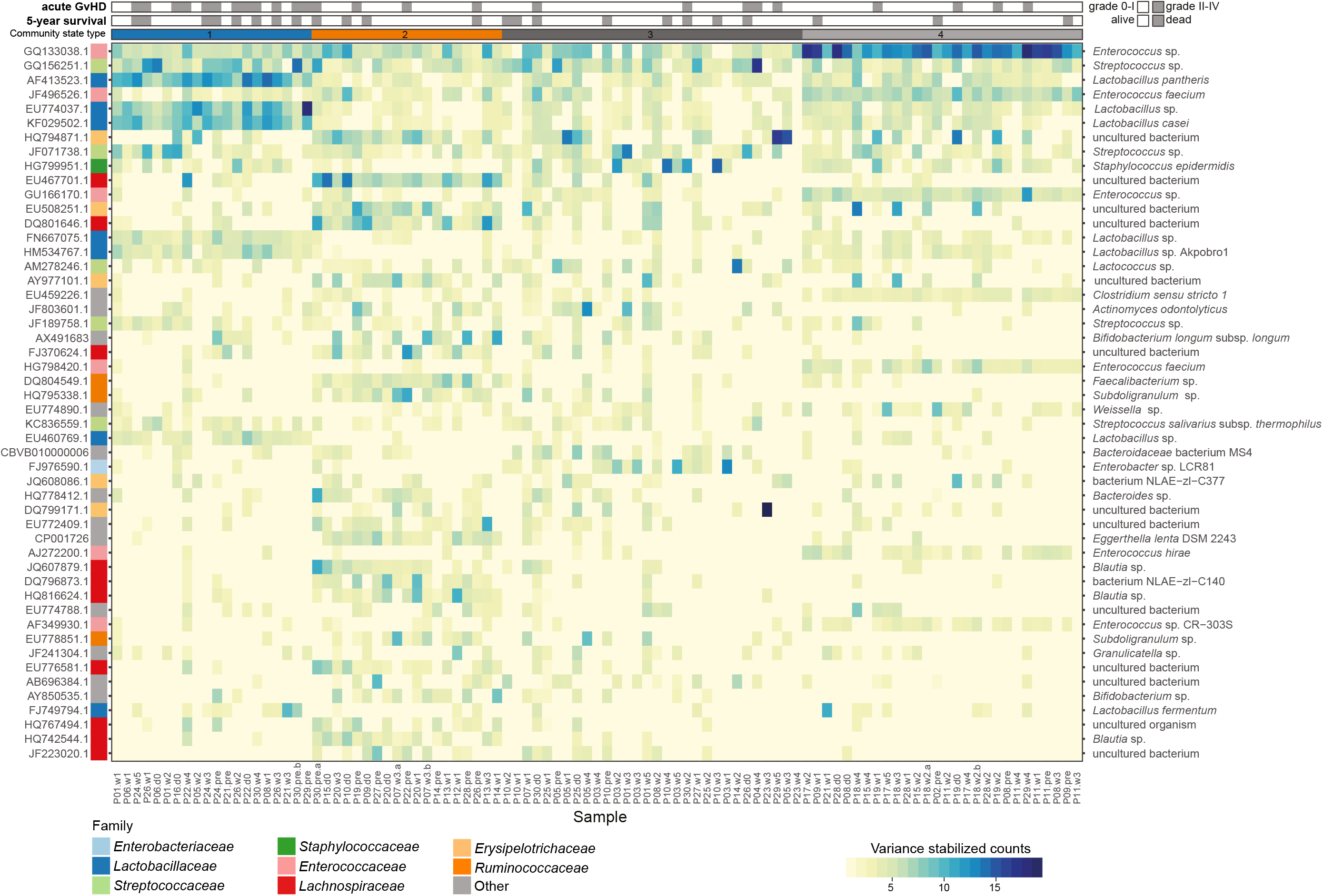
Community state types and gut microbial patterns. Heat map of variance stabilized counts of the 50 most abundant OTUs of the intestinal microbiome over all samples, grouped into community state types (CSTs). Based on their OTU-composition, samples were assigned to community state types (CSTs) by partitioning around medoid (PAM) clustering using Bray-Curtis distance. The optimal number of clusters (k = 4) was estimated from the gap statistic and Silhouette width validation. Members of the *Lactobacillaceae* family dominated the abundance profiles within CST 1. CST 2 exhibited domination by *Lachnospiraceae*, *Erysipelotrichaceae* and *Ruminococcaceae* members. *Enterobacteriaceae*, *Streptococcaceae* and *Staphylococcacea* were characteristic for CST 3. CST 4 was characterized by a high abundance of *Enterococcaceae*. Average Silhouette width was s(*i*) = 0.16 (range: −0.02 – 0.36), with CST 1 and CST 4 being the best defined clusters (s(*i*) = 0.23 and 0.36, respectively). A Silhouette coefficient s(*i*) close to 1 indicates appropriate clustering of the respective samples. Sample names at the bottom indicate patient (P) and time point (pre: pre-HSCT, d0: day of HSCT, w: week). An “a” or “b” indicates that two samples were collected from the respective patient at the same time point, but on two different days. (PDF)

**Additional file 9: Figure S6.**
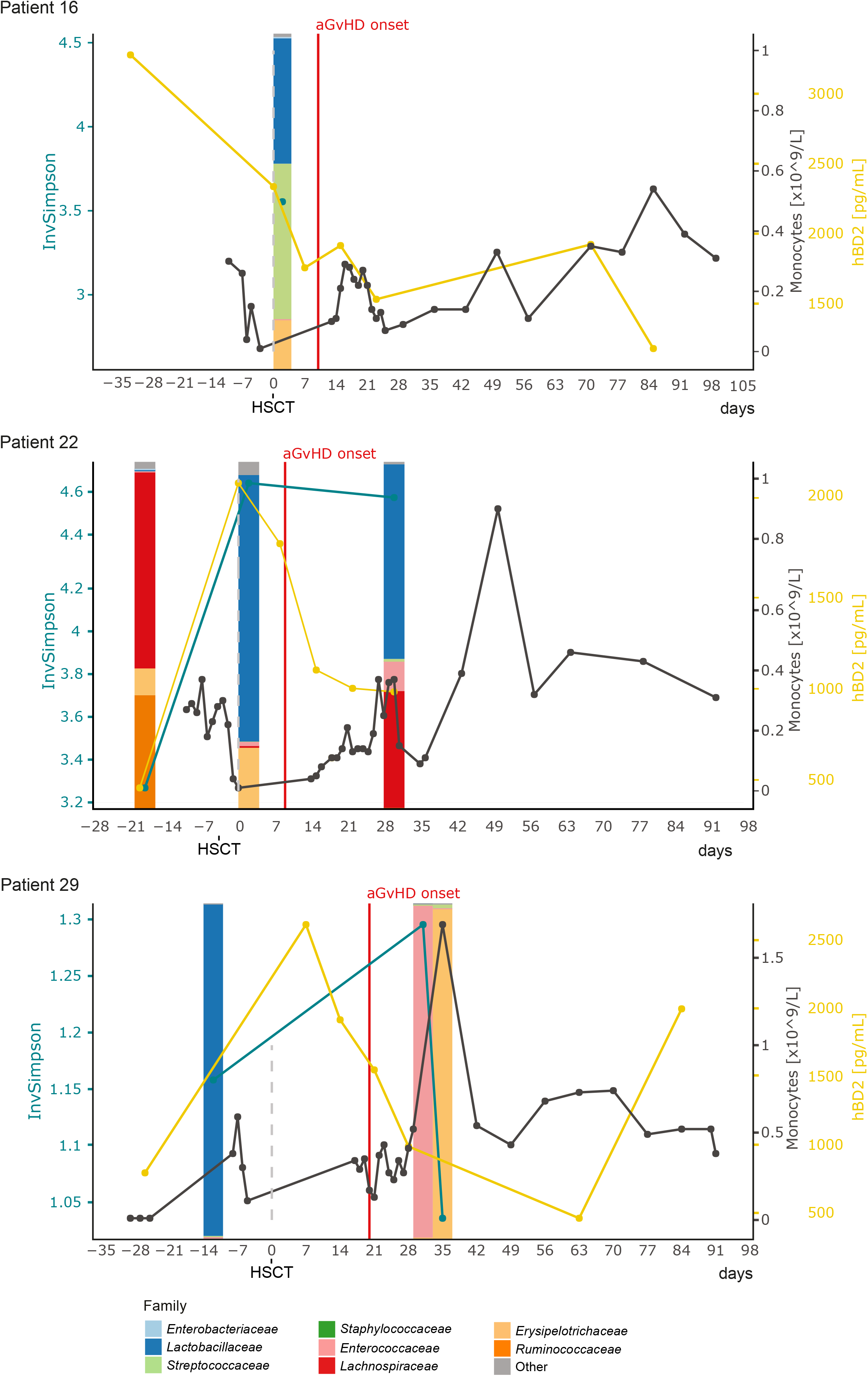
Longitudinal profiles of microbial community composition and immune markers in patients with aGvHD who survived. In three representative patients with moderate to severe aGvHD who survived, high abundance of *Lactobacillaceae* was observed already before aGvHD onset. None of the depicted patients had a bacterial infection recorded during the monitored period. InvSimpson, inverse Simpson diversity index; hBD2, human beta-defensin 2. (PDF)

Additional file 10: **Supplementary Discussion**. Discussion concerning survival following high *Lactobacillaceae* abundances prior to the onset of aGvHD, and associations of adverse outcomes with *Enterococcus* compared with previous studies. (PDF)

Additional file 11: **R data analysis report 1**. Data preparation, filtering and transformation. (HTML)

Additional file 12: **R data analysis report 2**. Bacterial alpha-diversity over time and rank abundance curve for the gut microbiome of HSCT patients. (HTML)

Additional file 13: **R data analysis report 3**. Temporal patterns of immune markers and immune cells in HSCT patients. (HTML)

Additional file 14: **R data analysis report 4**. Correlations between immune markers, immune cell counts, and outcomes in HSCT patients. (HTML)

Additional file 15: **R data analysis report 5**. Variable selection and multivariate analyses of immune parameters and intestinal bacterial taxa in HSCT patients. (HTML)

Additional file 16: **R data analysis report 6**. Clustering of samples into Community State Types (CSTs) based on Jenson-Shannon divergence. (HTML)

Additional file 17: **R data analysis report 7**. Longitudinal profiles of microbial community composition and immune markers. (HTML)

Our results suggest that a high *Lactobacillaceae* abundance prior to the onset of aGvHD may point to a preventive effect, as these patients survived. A human clinical trial of *Lactobacillus rhamnosus* GG prebiotic gavage to HSCT patients at time of engraftment demonstrated no protection against GvHD [1]. This could mean that *Lactobacillaceae* may not play a key role in aGvHD development, at least not the particular *Lactobacillus rhamnosus* strain under the conditions used in this group of patients. However, the administered probiotic did not alter the abundance of *Lactobacillus* spp. in the patients’ guts [1], suggesting that the strain was not able to establish and proliferate in the host environment in this situation. An intrinsic increase of *Lactobacillaceae* prior to aGvHD onset, as observed here, therefore might still play a role in reducing aGvHD. A recent study related to the use of a probiotic given to infants to prevent sepsis suggested that the time point of application of specific *Lactobacillus* sp. strain as a synbiotic played a critical role in positive clinical outcomes[2]. Furthermore, a study on gut microbial immunomodulation emphasized the importance of characterizing bacteria at the strain-level, because individual strains can have different modulatory effects on the immune system [3]. Therefore, it would be of great interest to determine the identity and predicted function of the specific *Lactobacillus* spp. strains in our patients, and in particular, in those who exhibited an early high abundance of *Lactobacillus* spp., as compared with those who experienced an expansion of *Lactobacillus* spp. after aGvHD and who later died.

Interestingly, *Enterococcus* was not among the most relevant taxa identified by our multivariate analyses. Intestinal domination of *Enterococcus* spp. was not clearly associated with adverse outcomes in our subgroup of 30 patients, in contrast to previous findings [4–6]. It should be noted that these previous observations were made in adult allo-HSCT patients and were dependent on the type and amount of antimicrobial treatment. In addition, to elucidate this discrepancy further, we are currently characterizing *Enterococcus* isolates from fecal samples of our patient group, to gain insight into bacterial strain-level differences.

